# Structural basis for nutrient-activated channel opening in bacterial spore germination receptors

**DOI:** 10.64898/2026.08.27.747515

**Authors:** Joshua C. Cofsky, Fernando H. Ramírez-Guadiana, Briana L. Sobecks-Doherty, Lior Artzi, Jeremy D. Amon, Yongqiang Gao, Ron O. Dror, Josefina del Mármol, David Z. Rudner, Andrew C. Kruse

## Abstract

GerA-family germinant receptors were recently identified as ion channels that initiate germination of dormant bacterial spores in response to nutrients. Here we report 2.0-Å cryo-electron microscopy structures of apo, agonist-bound, and antagonist-bound GerA, revealing a symmetric pentamer of heterotrimers. Signaling relies on two allosteric switches that radiate from the LeuT-type ligand-binding subunit in opposite directions, one away from the channel subunit and one directly toward it. A lipoprotein subunit reroutes the first switch and allosterically activates the second, promoting their convergence on the channel. Together, these rearrangements comprise a multipronged, cooperative signal transduction pathway that connects nutrient binding to pore opening in this new class of ligand-gated ion channels.

## INTRODUCTION

When starved of nutrients, diverse bacteria differentiate into metabolically inactive, sterilization-resistant spores that can remain viable for centuries (*1–3*). Dormant spores constantly monitor their environment for nutrient signals. Upon exposure to germinants, spores rapidly exit dormancy, returning to a vegetative state capable of growth, division, and, in many cases, infection. Thus, cycles of sporulation and germination are major drivers of antibiotic resistance and represent key targets in infectious disease research (*4*). Genetic studies over the last fifty years have identified a conserved family of nutrient-sensing receptors, typified by GerA from *Bacillus subtilis*, as the initiators of germination (*5–7*). We recently discovered that GerA-family receptors function as a new class of ligand-gated ion channels (*8*). However, GerA’s molecular structure, as well its mechanism of nutrient detection and signal transduction, remain unknown.

The GerA nutrient receptor complex, which is activated by L-alanine and a subset of other L-amino acids (*9–12*), contains three subunits: integral membrane proteins GerAA and GerAB, as well as lipoprotein GerAC (*7*). Previous work established GerAB’s homology to LeuT-like secondary active transporters and provided strong evidence that this subunit contains the receptor’s nutrient binding site (*10*, *13*). GerAA has no known homologs outside the germinant receptor family, but structure predictions and mutational studies have pinpointed this protein as the channel-forming subunit (*8*). GerAC, also without homologs, is essential for GerA function (*8*, *14*), but its role is unknown. It is unclear how these three subunits cooperate to transmit a germinant signal from GerAB’s ligand-binding pocket to GerAA’s channel pore.

Our current understanding of this signal transduction process comes from a genetic screen that identified signal-suppressing mutations in GerA (*15*). Interestingly, these mutations were scattered across the entire GerA complex, suggesting a complicated signaling network spanning the three protein subunits. However, most suppressive mutations were clustered in a GerAA α-helix (residues 310-320) that is predicted to lie perpendicular to the central channel axis, abutting the channel pore. Epistatic analysis indicated that this helix, which we term the “transducer helix,” functions late in the signaling pathway, perhaps serving as the final link in the allosteric chain connecting nutrient binding to pore opening.

Here, we report cryo-EM structures of GerA in apo, agonist-bound, and antagonist-bound states. These snapshots, combined with mutational analysis and structure predictions, revealed that two allosteric switches radiate from the GerAB ligand-binding site in opposite directions. Aided by GerAC, the two signals converge on GerAA’s transducer helix, the gatekeeper of channel opening. These findings establish how this ion channel complex converts information about environmental nutrient concentration into a developmental decision.

## RESULTS

### Generating well-ordered GerA receptor complexes for structural analysis

After multiple attempts to purify the intact GerA complex from *E. coli*, we developed a system for preparative-scale co-expression of FLAG-tagged GerAA, untagged GerAB, and untagged GerAC from three chromosomal loci in *B. subtilis* (see Methods). Detergent-solubilized GerA complexes, purified by FLAG affinity and size exclusion chromatography, were monodisperse (**Fig. S1A**) and contained all three subunits (**Fig. S1B**). Analysis of the sample by cryo-EM revealed a flower-shaped *C*_5_-symmetric complex, with a GerAA channel at the center and GerAB petals decorating the periphery (**Fig. S2A**).

While GerAC was full-length and tightly biochemically associated with the complex in this sample (**Fig. S1B**), only its lipidated N-terminus was visible in the cryo-EM map, sandwiched between GerAA and GerAB (**Fig. S2A**). Because many AlphaFold-predicted GerAC-GerAA and GerAC-GerAB interactions were missing from this map, we suspected GerAC’s disorder was an artifact of *in vitro* reconstitution. To stabilize these interactions, we engineered and purified a set of cysteine mutants that, based on AlphaFold modeling, were predicted to form inter-chain disulfide bonds (**Fig. S2B**). We chose two candidates (GerAA(A318C):GerAB:GerAC(P52C) and GerAA(A318C):GerAB:GerAC(S56C)) for further analysis based on their strong disulfide formation (**Fig. S2C**), GerAC density in negative-stain EM micrographs (**Fig. S2D**), and ability to support robust germination when expressed in spores (**Fig. S3**, **Supp. Note 1**). Cryo-EM imaging of the disulfide-linked complexes revealed successful GerAC ordering (**Fig. 1A**, **Fig. S4**). We focused our subsequent structural analyses on the disulfide-linked complexes, but comparisons to structures of the wild-type complex helped guide models for GerAC function (**Table S1**, **Fig. S4**).

**Fig. 1.**
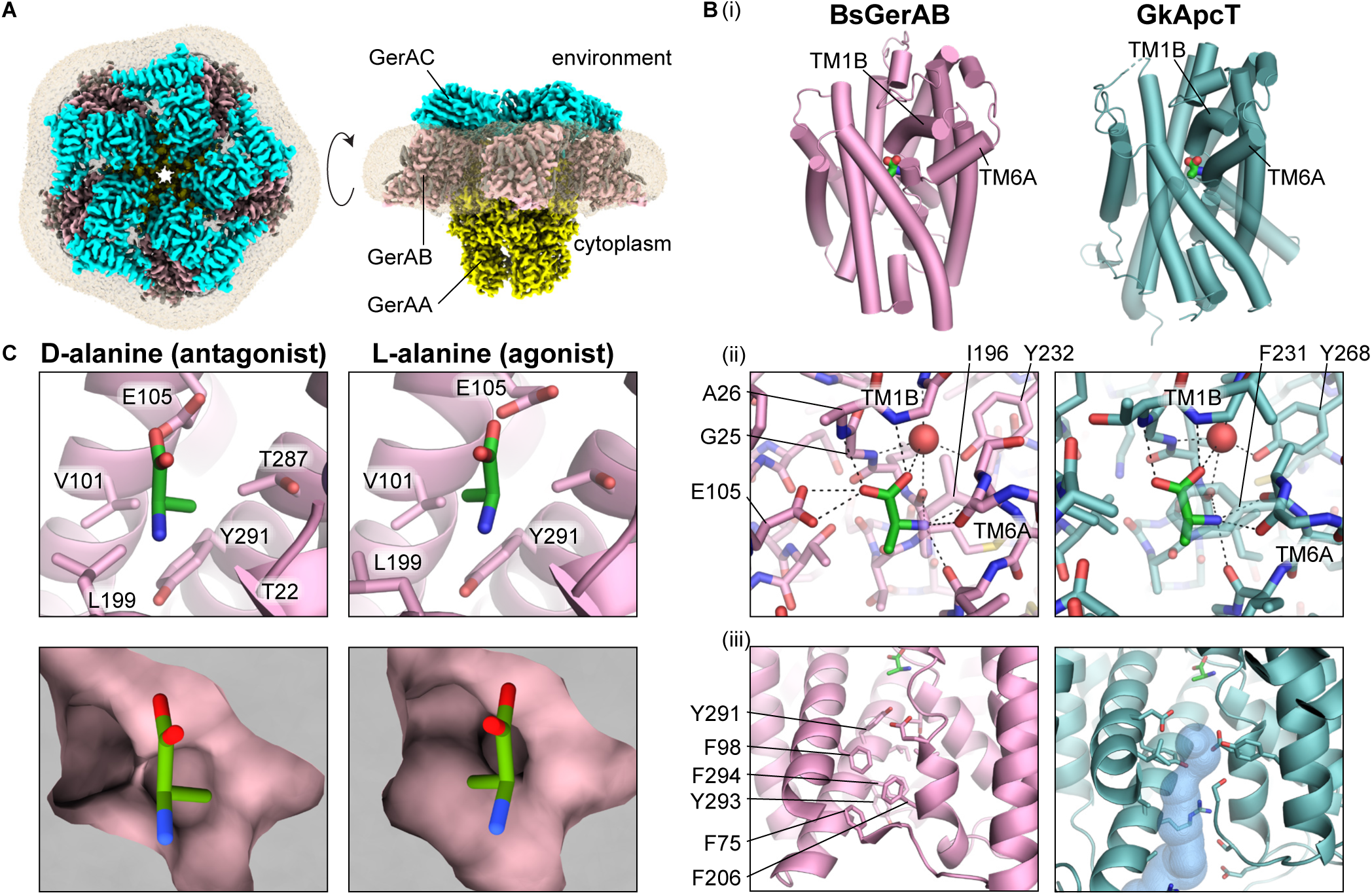
GerA nutrient receptor architecture and LeuT-like ligand-binding mechanism. (**A**) L-alanine-bound GerAA(A318C):GerAB:GerAC(S56C), unsharpened composite map. Ordered detergent, opaque gray. Disordered detergent, translucent beige (unsharpened consensus map). (**B**) (**i**) L-alanine-bound structures of *B. subtilis* GerAB (extracted from structure in (A)) and *G. kaustophilus* transporter ApcT (pdb_00005oqt). The ligand in the deposited ApcT model lacked a carboxylate oxygen; it was repaired by reference to the L-alanine in the GerAB model. (**ii**) Carboxylate- and amine-binding architecture. Ligand-capping sidechains GerAB-I196 and ApcT-F231 are translucent for clarity. Water molecule, red sphere. (**iii**) MOLE-calculated (*66*) cytoplasmic tunnel, blue surface. View-obstructing residues removed for clarity. (**C**) Sidechain-sensing cavity of D-alanine-bound and L-alanine-bound GerAB, within GerAA(A318C):GerAB:GerAC(S56C).

### GerAB binds amino acids using the canonical LeuT transporter mechanism

We first analyzed how GerA binds diverse amino acids and discriminates between them. L-alanine and L-valine activate GerA signaling while larger amino acids do not, and D-alanine competitively inhibits activation (*9*–*12*), suggesting a common binding mode for the amine and carboxylate but size- and stereochemistry-specific interactions around the sidechain. As predicted previously (*10*), L- and D-alanine are bound within the breakpoint of GerAB’s discontinuous TM1 helix using a mechanism resembling that of GerAB’s homolog, the LeuT-type transporter GkApcT (**Fig. 1Bi**) (*16*). The alanine amine is held by backbone carbonyls contributed by TM1 (GerAB-M23) and TM6 (GerAB-I196, S197, L199) (**Fig. 1Bii**). The alanine carboxylate is held by backbone amines (GerAB-A26 and G27) at the positively charged tip of TM1B, as well as a water molecule coordinated by the backbone at TM1’s helical break (GerAB-M23 to L28) and a conserved tyrosine (GerAB-Y232) (**Fig. 1Bii**). This amino acid binding mechanism explains the previous observation that GerAB(G25A) fails to signal in response to L-alanine (*10*), as a methyl sidechain at position 25 would sterically occlude the ligand alanine carboxylate and twist the TM1 backbone away from its helix-breaking trajectory via Ramachandran constraints.

There is one key contact that appears in structures of GerAB but not of GkApcT: a hydrogen-bonding interaction between the ligand alanine carboxylate and the GerAB-E105 carboxylate (**Fig. 1Bii**). Comparing the apo/D-alanine-bound structures to the L-alanine-bound structure, E105 undergoes a rotamer flip (**Fig. 1C**), which likely plays a role in signal transmission (see Switch 2 discussion). Support for the importance of E105 comes from previous functional analyses of the GerAB(E105K) mutation, which completely abrogated L-alanine-activated germination but allowed slow, alanine-independent germination when coupled with a GerAA hypermorph (*15*).

The final non-sidechain-specific feature of the ligand-binding mechanism is GerAB-I196, which resembles GkApcT’s outward-occluding cap residue F231 and likely inhibits amino acid dissociation (**Fig. 1Bii**) (*16*). Consistent with this role, spores containing GerAB(I196G) failed to respond to L-alanine (**Fig. S5**, **Supp. Note 1**) despite unperturbed GerA complex stability (**Fig. S6**).

Our cryo-EM structures revealed that GerAB discriminates between different amino acid sidechains using a spacious cavity lined by GerAB-V101, L199, and Y291, which we term the sidechain-sensing cavity (**Fig. 1C**). The L-alanine sidechain methyl points into this cavity but occupies only a fraction of its volume, explaining how the larger sidechain of L-valine is also accommodated. Importantly, we previously demonstrated that GerAB’s germinant profile could be broadened by making mutations to V101 and L199 that either expanded the cavity’s size or changed its chemical character (*10*).

The carboxylate and amine of the competitive inhibitor D-alanine are braced similarly to those of L-alanine. In this pose, D-alanine’s sidechain methyl points away from the cavity’s spacious center and instead toward its wall, where it contacts the backbone carbonyl of GerAB-T22 and the γ-methyl of GerAB-T287 (**Fig. 1C**). In support of these stereochemistry-specific contacts, relieving the methyl-methyl contact by mutating GerAB-T287 to serine improved D-alanine’s relative inhibitory potency (**Fig. S7**). These observations explain how the same pocket can bind either enantiomer: a shared tight binding mode for the carboxylate/amine and a more permissive cavity around the sidechain.

Despite the similarity between GerAB and GkApcT with regard to global protein fold and amino acid binding mechanism, the cytoplasmic tunnels of the two proteins are strikingly different. Whereas GkApcT contains polar sidechains to permit amino acid entry to the cytosol (*16*), GerAB contains tightly packed hydrophobic sidechains that occlude the transmembrane conduit (**Fig. 1Biii**). These findings suggest that during evolution, GerAB lost the inward-facing half of its ancestral transport function so that its outward-facing functions could be repurposed entirely as a signaling switch.

### Transmission of the nutrient-binding signal employs the outward-occluding helix of the LeuT transporter fold

We inspected the structures for ligand-dependent conformational differences that could reveal how information is transmitted from GerAB’s ligand-binding site to the GerAA channel, focusing first on GerAB-TM6A (**Fig. 2A**). In GerAB’s transporter cousins, TM6A functions in outward occlusion and thus undergoes large conformational changes while regulating solute entry and release (*16–20*). Similarly, in the process of grasping the amine of either L- or D-alanine, the backbone carbonyls at the base of TM6A (GerAB-I196 and S197) tug the entirety of TM6A towards the ligand. While the amine groups of both agonist and antagonist perform the same carbonyl-tugging action, the consequences for TM6A structure are strikingly different. In the transition from the apo structure to the D-alanine-bound structure, TM6A undergoes only minor bending toward the ligand without loss of helicity. By contrast, L-alanine causes the helix to unravel, most dramatically manifested in the transition of GerAB-V193, V194, and S195 toward a β-extended conformation (**Fig. 2A**). The distinct changes caused by the two ligands are likely due to their subtly different poses within the binding site: L-alanine tilts toward GerAA in a way that more dramatically distorts the base of TM6A (**Fig. 2A**, **Fig. S8**, **Supp. Note 2**). To test the importance of helix unraveling in GerA activation, we introduced helix-breaking prolines into TM6A. GerAB(V193P) and GerAB(V193P, V194P, S195P) each yielded an increase in the appearance of phase-dark spores at the end of sporulation, a hallmark of hyperactive germination (**Fig. 2B**, **Fig. S5**, **Supp. Note 1**). Additionally, when purified spores containing GerAB(V193P, V194P, S195P) were exposed to L-alanine, germination proceeded markedly faster than in the wild-type control spores (**Fig. S5**, **Supp. Note 1**). We conclude that nutrient-activated TM6A helix unraveling promotes GerA activation.

**Fig. 2.**
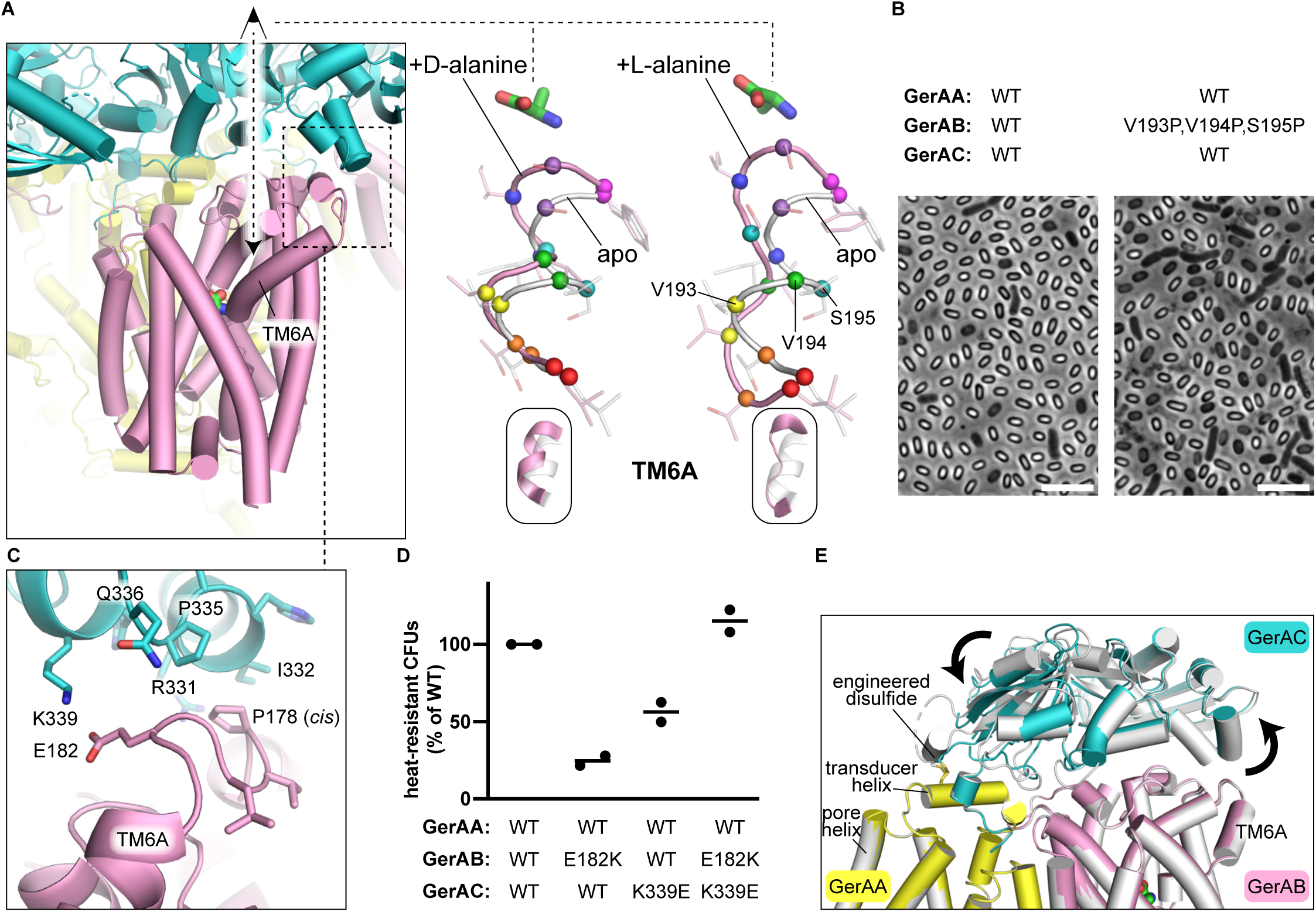
Mechanism for nutrient-mediated activation of Switch 1, pointing away from the GerAA channel. (**A**) GerAB-TM6A movements in response to D- and L-alanine. (**B**) Phase-contrast light micrographs of sporulating *B. subtilis* containing WT GerA or GerA with TM6A-unraveling mutations. Scale bars, 5 µm. (**C**) Peripheral GerAB-GerAC contact transducing GerAB-TM6A movements to GerAC; D-alanine-bound GerAA(A318C):GerAB:GerAC(S56C). (**D**) Quantification of heat-resistant spore formation by *B. subtilis* containing mutations at the peripheral GerAB-GerAC contact. (**E**) Comparison of L-alanine-bound GerAA(A318C):GerAB:GerAC(S56C) (white) to L-alanine-bound GerAA(A318C):GerAB:GerAC(P52C) (colored), demonstrating the “seesaw” motion of GerAC and its effect on GerAB-TM6A.

Interestingly, TM6A lies on the periphery of the GerA complex, indicating that the signal initially propagates outward, away from GerAA’s pore. However, the top of TM6A interacts with the periphery of GerAC (**Fig. 2C**), suggesting that the lipoprotein could redirect the signal. We investigated whether the peripheral GerAB-GerAC interaction was important for GerA function by mutating key interacting residues and measuring the effect on sporulation and germination. GerAB(P178V) and GerAC(R331A, I332A, P335A, Q336A) each resulted in loss of GerAA in spores (**Fig. S6**), indicating that the structural elements at this interface are crucial for the stability of either individual subunits or the intersubunit interaction (**Supp. Note 1**). Furthermore, GerAB(E182K), GerAC(K339E), and GerAC(I332A) each caused a decrease in post-sporulation CFU counts (**Fig. 2D**, **Fig. S5**) without altering GerAA levels (**Fig. S6**), consistent with impaired GerA signaling. Notably, GerAB-E182 and GerAC-K339 form a salt bridge (**Fig. 2C**). When the GerAB(E182K) and GerAC(K339E) mutations were combined to effect a charge swap, normal germination behavior was restored (**Fig. 2D**, **Fig. S5**). These findings establish the functional importance of the peripheral GerAB-GerAC interaction.

We propose that, during GerA activation, ligand-induced unraveling of the base of GerAB’s TM6A shifts the top of the helix, which in turn pushes on GerAC. GerAC spans the entirety of the complex, contacting both GerAB’s peripheral TM6A helix and GerAA’s transducer helix previously implicated in pore opening (**Fig. 2E**). Thus, rigid-body motions of GerAC would couple peripheral structural changes to central ones. Evidence for this coupling function comes from comparison of cryo-EM structures containing the two different GerAA-GerAC disulfide linkages. The GerAA(A318C)-GerAC(P52C) disulfide pulls GerAC further down in the center than the GerAA(A318C)-GerAC(S56C) disulfide. Like a seesaw, the periphery of GerAC rises in response, bringing the top of GerAB TM6A with it (**Fig. 2E**). While our cryo-EM structures of L-alanine-bound GerA lack major ligand-induced changes to GerAC position, these structures likely represent a pre-activated state (see Discussion), and we propose that more dramatic changes to GerAC position occur during full activation. We assign the term “Switch 1” to the combination of nutrient-activated GerAB-TM6A helix unraveling and GerAC’s redirection of the signal toward the GerAA channel.

### A second switch transmits the germinant signal directly from GerAB to GerAA with allosteric enhancement by GerAC

We next investigated ligand-induced structural changes between the GerAB ligand-binding site and the GerAA channel, opposite Switch 1 (**Fig. 3**, **Supp. Note 3**, **Fig. S9**). L-alanine binding causes a rotation of GerAB-TM8 that extends from F284, in plane with the ligand (**Fig. 3A**), to E273, at the top of the helix (**Fig. 3B**). Importantly, the top of TM8’s rotated segment presses against the peripheral tip of GerAA’s transducer helix (**Fig. 3A-B**, **Supp. Note 3**, **Fig. S9**), providing a mechanism by which TM8 conformation could influence GerAA channel opening. We assign the term “Switch 2” to the nutrient-activated TM8 rearrangement.

**Fig. 3.**
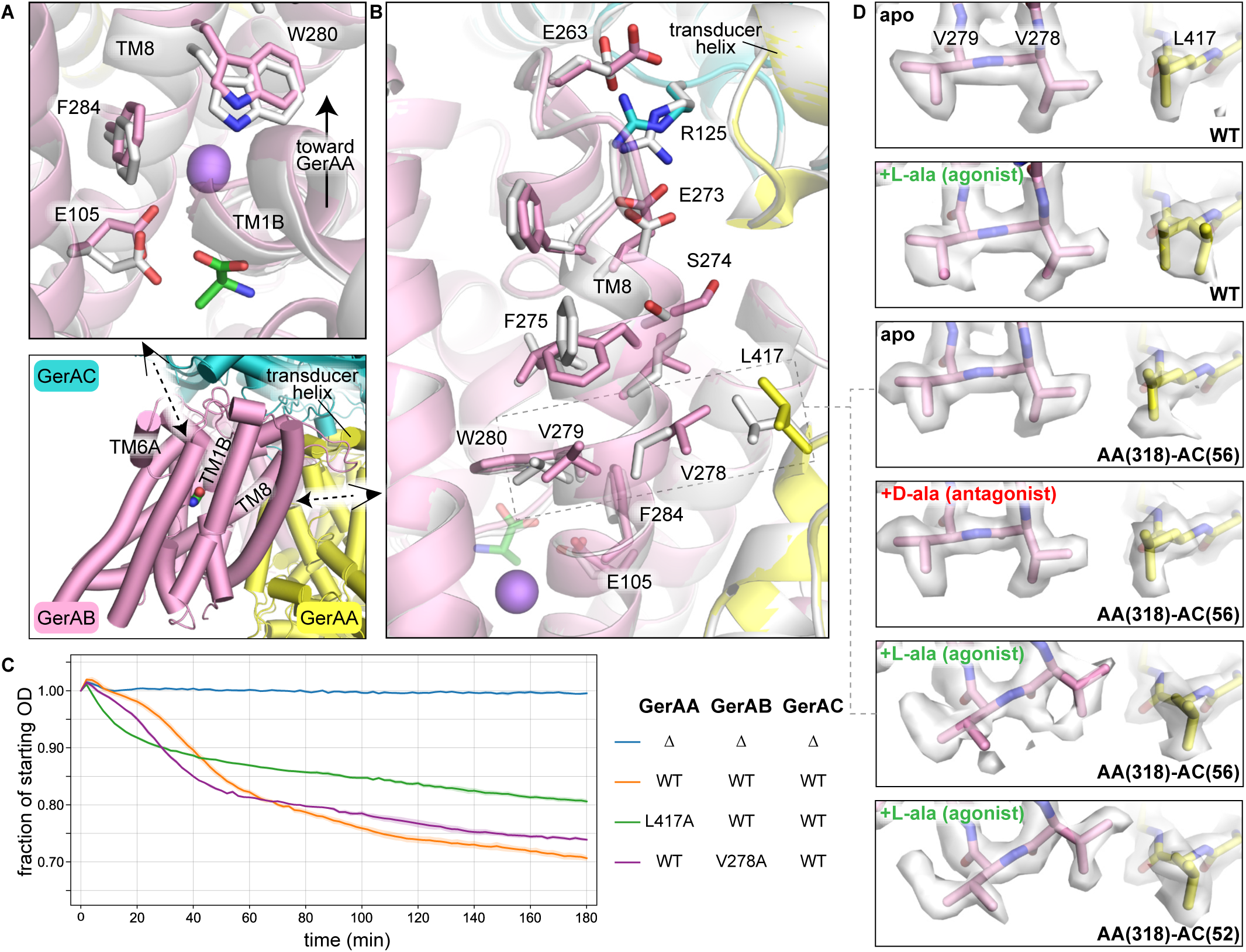
Mechanism for nutrient-mediated activation of Switch 2, pointing toward the GerAA channel. (**A**) Detail of Switch 2 near L-alanine (green). Comparison of apo (white) and L-alanine-bound (colored) GerAA(A318C):GerAB:GerAC(S56C). Na^+^ ion, purple sphere. (**B**) TM8 helix rotation that comprises Switch 2, activated by L-alanine at the bottom, pressing against GerAA transducer helix at the top. (**C**) Germination kinetics of spores with mutations in Switch 2’s GerAA/GerAB interface. Shaded area, standard error of the mean for three technical replicates. See **Supp. Note 1** for details of assay interpretation. (**D**) Cryo-EM maps (consensus, sharpened) and models demonstrating ligand- and GerAC-dependence of Switch 2 activation. Both rotamers of GerAA-L417 were modeled for L-alanine-bound WT GerA due to comparable occupancy. For an explanation of AA(318)-AC(52)’s greater occupancy of the activated state than AA(318)-AC(56), see **Supp. Note 3**.

We examined the structural features controlling Switch 2 propagation, starting at the base of TM8’s rotated segment. The rotation originates with the motion of several ligand-proximal residues toward GerAA, including a rotamer flip of GerAB-E105 and a global movement of helix TM1B (**Fig. 3A**). These residues interact at a TM1-TM8 interface spanned by an octahedrally coordinated Na^+^ ion (**Fig. 3A**), which could explain previous observations of salt sensation by GerA and its homologs (*21–27*). Mutation of TM8 residues near this interface (GerAB-F284, W280, or V279) resulted in either impairment or complete loss of germination, suggesting that TM8 rotation is crucial for germination activation (**Fig. S5**).

Halfway up TM8’s rotated segment, GerAB-V278 contacts GerAA-L417, causing GerAA-L417’s rotamer to flip in the presence of L-alanine (**Fig. 3B**). Mutation of either GerAB-V278 or GerAA-L417 to alanine resulted in fewer heat-resistant spores, more phase-dark spores, and faster germination kinetics, indicating that the GerAB-V278/GerAA-L417 interaction plays an autoinhibitory role (**Fig. 3C**, **Fig. S5**). Notably, the segment of the GerAA-TM1 helix that contacts GerAA-L417 is a known site of variation between common *B. subtilis* lab strains with different germination phenotypes (*28*), suggesting that GerAA-TM1 influences this autoinhibitory interface.

At the top of GerAB-TM8’s rotated segment, GerAC-R125 pulls on GerAB-E273 in a manner that appears to stabilize the TM8 rotation, suggesting that GerAC potentiates Switch 2 propagation (**Fig. 3B**). Consistent with this hypothesis, mutation of GerAB-E273 or GerAC-R125 severely impaired germination (**Fig. S5**, **Supp. Note 3**). We further investigated the hypothesis by comparing the cryo-EM maps for complexes with disordered (WT) vs. ordered (disulfide-stabilized) GerAC, which revealed GerAC-dependent changes in the relative occupancy of the two observed Switch 2 conformations. Without GerAC ordering, Switch 2’s activated conformation is intermediately populated near the ligand (**Fig. S10A**), sharply attenuated at the GerAB-V278/GerAA-L417 midpoint (**Fig. 3D**), and undetectable at the top of TM8 (**Fig. S10B**). We conclude that GerAC is an allosteric enhancer of Switch 2 that allows the ligand-binding signal to propagate all the way to GerAA’s transducer helix.

### GerAA’s pore helices are mobile but maintain a closed channel in the cryo-EM structures

The pore helices (GerAA-TM3) appear essentially identical in all cryo-EM maps, irrespective of GerAC ordering or GerAB ligand binding (**Fig. S11A**, **Table S2**). This uncoupling of ligand binding from pore transitions is a common result for structural studies of ion channels, where *in vitro* sample stabilization techniques make rare open-pore conformations difficult to capture (*29*). In all maps, the five GerAA-V362 sidechains are well resolved, pointing into the lumen in a symmetric configuration with a pore diameter of <4.1 Å (**Fig. S11B**). A hydrophobic constriction of this size suggests a non-conductive channel (*30*). As further evidence for this residue’s key role in pore constriction, mutation of V362 to alanine causes phenotypes consistent with a constitutively open state, while mutation to leucine causes permanent channel closure (*8*).

Interestingly, in contrast to GerAA-V362, the other pore helix sidechains are poorly resolved in the cryo-EM maps (**Fig. S11C**). For several of these residues, mutation causes strong germination phenotypes (*8*), suggesting that they, like V362, also play important channel gating roles. To probe channel structure along the entire pore, we performed molecular dynamics simulations of the GerAA transmembrane domain and monitored the constrictions formed by all pore-facing sidechains. Across 12 independent simulations, each 2 µs in length, the channel remained closed due to constrictions formed by Q354, L358, V362, and Q366, despite some flexibility at these residues (**Fig. 4A**, **Fig. S12**). We conclude that the channel in the cryo-EM sample is in a closed conformation.

**Fig. 4.**
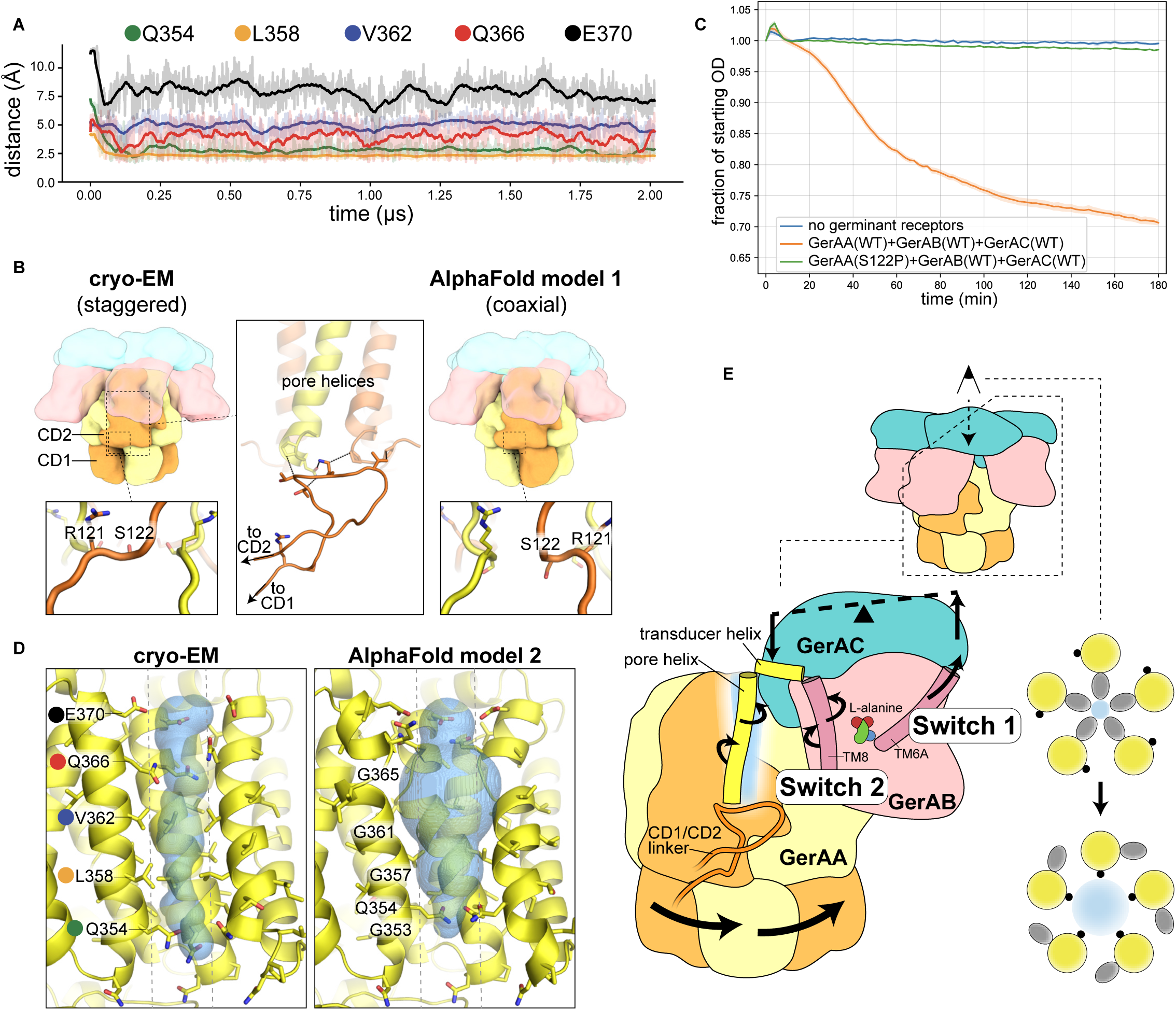
Structural details of GerAA’s pore helices/cytoplasmic domains and overview mechanistic model. (**A**) Molecular dynamics simulation of the GerAA transmembrane domain. “Distance” is the minimal pairwise distance between residues from non-adjacent protomers, which estimates pore size (see Methods). For reference, the diameter of a hydrated K^+^ ion is 6.6 Å (*67*, *68*). (**B**) Comparison of cryo-EM model of L-alanine-bound GerAA(A318C):GerAB:GerAC(S56C) and AlphaFold model of pentameric GerA (15-Å lowpass filter). Coloring of GerAA protomers alternates between yellow and orange to indicate which CD1 is linked to which CD2. Large inset: trajectory of the GerAA-CD1/CD2 linker and its contact with the pore helices. See **Fig. S14A** for detail. Small insets: linker residues whose backbone dihedrals underlie the staggered vs. coaxial CD1 positioning. (**C**) Germination kinetics of spores with mutation in the CD1/CD2 linker. Shaded area, standard error of the mean for three technical replicates. (**D**) GerAA channel pore of cryo-EM model (L-alanine-bound GerAA(A318C):GerAB:GerAC(S56C)) and AlphaFold model (different model from that shown in (B), see **Table S2**). Two protomers are removed for clarity. MOLE-calculated (*66*) channel lumen (hydrogens excluded from calculation), blue surface. Dashed lines mark the edges of a cylinder around the pore axis with the same diameter as a hydrated K^+^ ion. (**E**) Left, overview model for nutrient activated GerA channel opening. In Switch 1, L-alanine binding causes GerAB-TM6A unraveling, a motion which GerAC transmits to the GerAA transducer helix via a “seesaw” action. In Switch 2, L-alanine binding causes GerAB-TM8 rotation, which presses directly on the GerAA transducer helix. The transducer helix tugs on the top of the pore helix, and the CD1/CD2 linker, tracking with the CD1 swivel, follows movements in the base of the pore helix. Right, cross-section of GerAA pore (blue), pore helices (yellow circles), bulky sidechains (gray ovals), and glycines (black circles). When the pore helices rotate in place, the bulky sidechain obstruction is removed and the channel opens.

### AlphaFold suggests new features of channel activation

Finally, we used AlphaFold (*31*, *32*) to generate hypotheses about features of the GerA activation mechanism that may not have been captured by our cryo-EM structures. While the five standard AlphaFold models were not trained to sample conformational ensembles, it was previously demonstrated that, when the predictions calculated by these models differ from each other or from experimental structures, they sometimes reflect alternative, biologically relevant conformations (*33–35*). Considering our cryo-EM structures alongside the predictions of the five AlphaFold models, we identified three key points of variation. The first difference, GerAC docking, is addressed in the supplementary material (**Supp. Note 4**, **Fig. S13**).

The second difference concerns cytoplasmic domains 1 and 2 of GerAA (GerAA-CD1/2), whose important regulatory role was identified in previous genetic studies but not mechanistically explained (*15*). Different AlphaFold models predict two different conformations for the pentameric ring of GerAA-CD1: one in which each CD1 is coaxial with its linked CD2 and another in which the domains are staggered (**Fig. 4B**, **Table S2**). The closed-pore cryo-EM structures all occupy the staggered conformation (**Fig. 4B**, **Table S2**). Because the long CD1/CD2 linker (GerAA residues 118-138) directly contacts the base of the pore helices (**Fig. 4B**, **Fig. S14A**) and because domain swapping affects gating in ionotropic glutamate receptors (*36*), we hypothesized that swiveling from the staggered to the coaxial conformation could be part of the channel-opening transition. The proposed swiveling action transfers each CD1 into the exact position previously occupied by its neighbor, which is a completely rigid-body change except for within a small region (GerAA residues 121-124) of the CD1/CD2 linker (**Fig. 4B**). We constrained the dihedral angle landscape of this linker region by making the GerA(S122P) mutation, which should bias the complex toward the staggered conformation and disfavor access to the coaxial conformation (**Fig. S14B**). Spores containing this mutation were unable to germinate (**Fig. 4C**, **Fig. S5**), despite maintenance of GerA stability (**Fig. S6**). These data argue that the GerAA-CD1 swiveling motion is required for channel opening.

The third difference concerns the pore helices. While some AlphaFold predictions of the GerAA channel resemble the closed cryo-EM structures, others exhibit varying degrees of change to the pore helix (**Table S2**). In one prediction, the central section of each pore helix (Q354 to G365) has ratcheted around its own helical axis by ∼90°, such that the position previously occupied by residue *n* is now occupied by its *n*-1 neighbor (**Fig. 4D**).

Notably, for all four gating residues (Q354, L358, V362, Q366), the *n*-1 neighbor is a glycine. In the rotated conformation, with the bulky sidechains replaced by glycines, only the Q354 sidechain still blocks the pore (**Fig. 4D**). We propose that GerAA channel opening involves pore helix rotation, though full channel opening may require additional rearrangements.

## DISCUSSION

Our structures reveal that the signal sent from GerA’s nutrient-binding site to its ion-conducting pore does not travel along a single direct path but rather along a branching web of interconnected conformational changes (**Fig. 4E**). Consistent with this idea, when a mildly germination-impairing Switch 1 mutation was combined with a mild Switch 2 mutation, the resulting impairment was synergistic (**Fig. S15, Fig. S5**). Thus, pore opening requires combined stimulation by multiple allosteric networks.

We speculate that this complexity evolved to fulfill a need unique to germinant receptors. Unlike the ligand-gated ion channels that underlie neuronal signaling, which open and close repeatedly on a millisecond timescale (*37*), germinant receptors are under immense evolutionary pressure to remain closed for years during periods unsuitable for vegetative growth, when even a single transient activation event would lead to cell death. Biological systems with low tolerance for inappropriate activation often leverage signaling cooperativity to effect noise suppression (*38*). Switch-like cooperativity emerges from a combination of oligomeric architecture (*39*) and the coupling energy contributed by interdependent structural transitions (*40*, *41*). Here, we identified both ingredients: we determined that germinant receptors have a stoichiometry of five, and we discovered several conformational rearrangements that likely energetically influence each other both within and across heterotrimeric subunits (**Fig. 4E**). Intriguingly, in spores, germinant receptors form clusters (*42*) in which inter-receptor contacts could further enhance signaling response sharpness (*43*), decreasing the frequency of spurious flickerings and unwanted exit from dormancy.

In previously characterized ionotropic glutamate receptors (iGluRs) and pentameric ligand-gated ion channels (pLGICs), which are not related to GerA, the ligand is bound by an extracellular soluble domain that perches atop the pore helices, allowing streamlined signal transmission (*44*, *45*). In germinant receptors, the germinant-binding pocket is housed in an integral membrane protein coplanar with the channel, requiring that the signal travel a long distance (∼60 Å from L-alanine C_α_ to pore center) via a circuitous, lipoprotein-facilitated path rather than by direct pore contact. To explain this complexity, we propose that evolution worked with the available proteome. For example, the ligand-binding domains of eukaryotic iGluRs likely arose from the soluble periplasmic binding proteins of prokaryotic ABC transporters (*46*, *47*). In *B. subtilis*, L-alanine import has been attributed to several standalone LeuT-like transporters (*48–51*) but none that employ a soluble partner protein for substrate delivery (*51*, *52*). Perhaps, in the organism that housed GerA’s evolutionary birth, the repertoire of repurposable L-alanine-binding proteins was similar (L-alanine, for unknown reasons, is the most common germinant across sporulating bacteria (*53*, *54*)). These repurposings raise the exciting possibility that an unappreciated diversity of genes annotated as transporters in genomic databases may instead encode signaling complex sensors.

Analogous evolutionary trajectories have already been identified in mammals. In the Kir6/SUR complex, the transmembrane component of a retired ABC transporter (SUR) was commandeered to regulate channel conductance (Kir6) in response to nucleotide concentration (*55*). In the mTORC1 pathway (which, like GerA, translates nutrient availability into cell growth decisions (*56–58*)), the LeuT-like arginine transporter SLC38A9 was co-opted for a sensing role. SLC38A9 has an auxiliary protein tail that, by default, is docked in the transport conduit but, upon competition by rising arginine levels, is released into the solvent where it can activate downstream signaling GTPases (*59*, *60*). Compared to SLC38A9’s simple sequestration-based mechanism, GerAB’s use of the same protein fold is quite sophisticated, involving a branching allosteric network that couples the motions of its transporter ancestor to rotation of a distant helix in a ligand-dependent manner.

In our cryo-EM structures, the nutrient-dependent conformational change of GerAA-L417 provides clear evidence for the arrival of ligand-binding information at the channel subunit, but the closed pore suggests that this conformation represents a pre-activated or desensitized state. Although our cryo-EM structures captured many crucial features of GerA’s activation pathway, it may be that full activation can only occur in the unusual desiccated environment of the inner spore membrane, characterized by high solute concentrations, large compressive forces, and immobilized macromolecules (*61–64*). Importantly, while AlphaFold provided hints as to what additional rearrangements accompany full activation, it failed to identify patterns in the coupling of these rearrangements (**Table S2**), highlighting the essential role of experimental structure determination in deciphering the functional meaning of conformational changes.

Many questions about GerA mechanism remain to be answered, most notably which ions it conducts and how their passage stimulates the release of dipicolinic acid by the SpoVA transporter (*65*). The abnormal environment of the inner spore membrane suggests that answers to some questions await challenging technical advances in the direct observation of these receptors functioning in their native clusters. Our high-resolution structures of the GerA complex, along with future mechanistic insights, may enable the development of new strategies to combat pathogenic bacteria that evade existing antibiotics by differentiating into sterilization-resistant spores (*4*).

## Supporting information

Supplementary Material Main PDF

## Acknowledgments

We thank all members of the Kruse, Rudner, del Mármol, and Dror labs for helpful advice, discussion, and encouragement. Negative-stain EM data were collected at the Harvard Medical School Electron Microscopy Facility. Preliminary cryo-EM data were collected at the UMass Chan Medical School Cryo-EM Core Facility and the UCSF Cryo-EM Core Facility, which is partially supported by NIH grants (S10OD020054, S10OD021741 and S10OD026881) and the Howard Hughes Medical Institute. Final cryo-EM data were collected at the Harvard Cryo-EM Center for Structural Biology at Harvard Medical School.

## Author contributions

Conceptualization: JCC, DZR, ACK

Funding acquisition: ROD, DZR, ACK

Investigation: JCC, FHR, BLS

Supervision: ROD, JdM, DZR, ACK

Visualization: JCC, BLS

Writing — original draft: JCC

Writing — review & editing: JCC, FHR, BLS, LA, JDA, YG, ROD, JdM, DZR, ACK

## Data, code, and materials availability

PDB and EMDB accession codes can be found in **Table S1**.

