## Supplementary Material Main PDF for "Structural basis for nutrient-activated channel opening in bacterial spore germination receptors"

### METHODS

GerA expression strain design

GerA subunits were co-expressed from a *Bacillus subtilis* 168 expression system developed for this purpose. We describe the strain used to produce WT GerA, whose genotype is Δ*ydiS*::lox72 Δ*gerA*::lox72 *yyzG*::*lacI*(lox72) *lacA*::$P_{veg}$-*ULP1*(lox72) *yvbJ*::$P_{hyperspank}$-MBP-SUMO-FLAG-*gerAA* (*erm*) *amyE*:: $P_{hyperspank}$-optRBS-*gerAB*(*cat*) *yhdG*:: $P_{hyperspank}$-optRBS-*gerAC*(*kan*). All GerA variants were based on WT. The restriction-enzyme-encoding *ydiS* gene was deleted to improve transformation efficiency during strain construction (*69*). The native *gerA* operon was deleted to avoid unwanted recombination during strain construction. A constitutively expressed *lacI* gene from *E. coli* was inserted at the *yyzG* locus and enabled isopropyl-β-D-1-thiogalactopyranoside-based control of the $P_{hyperspank}$ promoters, which drove robust expression of *gerAA*, *gerAB*, and *gerAC* inserted at the indicated chromosomal loci when paired with a ribosome-binding site optimized for strong translation initiation. BsmBI and BsaI restriction sites within the *gerA* genes were recoded to enable Golden Gate cloning. The 3' end of *gerAA*, which overlaps with the 5' end of *gerAB*, was also recoded. GerAB and GerAC had no tags. GerAA's N-terminal tag began with the first 15 nucleotides of the highly-expressed *B. subtilis metE* gene (encoding `MTTIK`), inspired by (*70*), followed by the maltose-binding protein gene from *E. coli*. Together, these tags improved expression. Finally, the SUMO-FLAG cassette enabled affinity purification. *In vivo* cleavage of this cassette by the SUMO-specific Ulp1 protease from *Saccharomyces cerevisiae* (expressed from the *lacA* locus under the control of a constitutive *B. subtilis* promoter) removed the MBP-SUMO and exposed the N-terminus of the FLAG tag, which is required for binding by the M1 α-FLAG antibody.

To predict where engineered cysteine mutations could stabilize GerAC, Disulfide by Design 2.0 (*71*) was run on all five models produced by the 5GerAA:5GerAB:5GerAC AlphaFold prediction. The six presented disulfides were the only ones detected by the program (default settings).

AlphaFold predictions

Using Colabfold 1.5.2 (*72*), AlphaFold2-Multimer (*31*, *32*) was run with default settings (with multiple sequence alignments) on either five copies of GerAA ("5GerAA") or five copies of all subunits ("5GerAA:5GerAB:5GerAC"), followed by addition of hydrogens and relaxation with AMBER. The signal peptide of GerAC was omitted. "AlphaFold model 1" (**Fig. 4B**, **Fig. S13**) is the top-ranked model from the 5GerAA:5GerAB:5GerAC prediction. "AlphaFold model 2" (**Fig. 4D**) is the second-ranked model from the 5GerAA prediction. See **Table S2** for a catalog of each AlphaFold model's conformational features.

GerA expression and purification

*B. subtilis* strains similar to that described above were inoculated into Terrific Broth (12 g/L Bacto Tryptone (gibco, #211705); 24 g/L yeast extract (VWR, J850); 0.4% (v/v) glycerol; 12.54 g/L $K_2HPO_4$; 2.31 g/L $KH_2PO_4$; 2 mM $MgCl_2$; 0.1% (w/v) glucose; 0.01% (v/v) Antifoam 204 (Sigma-Aldrich A6426)) and shaken at 37 °C until they reached $OD_{600}$ 0.025, at which point 25 µM IPTG was added. The culture was shaken at 37 °C for ~24 hours, at which point pellets were harvested and washed with HBS (20 mM Na-HEPES, pH 7.5; 150 NaCl).

Pellets were immersion-blended and Dounce-homogenized in lysis buffer (50 mM Na-HEPES, pH 7.5; 150 mM NaCl; 2 mM $MgCl_2$; 10% (v/v) glycerol; 0.0025 U/µL benzonase (Millipore E1014); 2 mg/mL chicken egg white lysozyme (Sigma-Aldrich L6876); 1X protease inhibitors). 1X protease inhibitors is: 1 mM phenylmethylsulfonyl fluoride; 1 µg/mL pepstatin A; 150 nM aprotinin; 10 µM leupeptin. Cell suspensions were stirred at 37°C for 1 hour to allow complete cell wall degradation, followed by another round of Dounce homogenization on ice. Lysed cells were centrifuged (168k x *g*, 70 min, 4 °C). Membrane pellets were resuspended in solubilization buffer (50 mM Na-HEPES, pH 7.5; 30% (v/v) glycerol; 1X protease inhibitors; 150 mM NaCl; 1% (w/v) sol-grade n-dodecyl-β-D-maltoside; 0.05 U/µL benzonase; 2 mM $MgCl_2$; 2 mM $CaCl_2$) using immersion blending and Dounce homogenization. These samples were rolled (2 hours, 4 °C) and centrifuged (168k x *g*, 70 min, 4°C), followed by supernatant collection. M1 α-FLAG resin was added to filtered supernatant, which was stirred overnight at 4 °C.

Resin was collected and washed with wash buffer 1 (20 mM Na-HEPES, pH 7.5; 10% (v/v) glycerol; 1X protease inhibitors; 150 mM NaCl; 2 mM $CaCl_2$; 0.05% (w/v) sol-grade DDM; 0.1% (w/v) glyco-diosgenin), then wash buffer 2 (remove DDM), then wash buffer 3 (reduce [GDN] to 0.02% (w/v)). The resin bed was resuspended in nuclease solution (wash buffer 3 plus: 0.5 U/µL benzonase; 2 mM $MgCl_2$; 2 U/µL nuclease P1 (NEB M0660S); 2 µM $ZnCl_2$) and allowed to digest overnight at 4 °C.

Resin was washed with wash buffer 3, followed by 1.5 hours of elution with elution buffer (20 mM Na-HEPES, pH 7.5; 10% (v/v) glycerol; 1X protease inhibitors; 150 mM NaCl; 5 mM ethylenediaminetetraacetic acid; 1 mg/mL FLAG peptide; 0.02% (w/v) GDN). For GerA complexes lacking a disulfide cross-link, 1 mM tris(2-carboxyethyl)phosphine was added to the affinity resin eluate and included in buffers at all downstream steps of purification, unless indicated otherwise. Eluate was concentrated (100-kDa molecular weight cutoff) to 400 µL, and 600 µL of 1.67X cryoprotectant buffer (20 mM Na-HEPES, pH 7.5; 150 mM NaCl; 0.02% (w/v) GDN; 75.5% (v/v) glycerol) was added, followed by flash-freezing and storage at −80 °C. Before any subsequent experiment, sample aliquots were thawed and purified by size exclusion chromatography on a Superose 6 Increase 10/300 GL column (Cytiva) in size exclusion buffer (20 mM Na-HEPES, pH 7.5; 150 mM NaCl; 0.02% (w/v) GDN). Standards for size exclusion chromatography were Bio-Rad #1511901. Standards for SDS-PAGE analysis were Bio-Rad #1610374. Purifying from 12 L of bacterial culture typically yielded ~1.7 mg of GerA complex in grid-ready format.

<u>Measuring disulfide completeness and strength</u>

Disulfide characterization was performed similarly to (*73*). GerA samples were size-exclusion-purified (TCEP was excluded for all protein variants) and concentrated. To initiate disulfide competition, protein sample (7.5 µL, $A_{280}$=2) was mixed with size exclusion buffer containing dithiothreitol (7.5 µL, 2X final concentration) and incubated at room temperature for 30 minutes. Disulfide exchange was quenched with 3 µL of size exclusion buffer containing 20 mM S-methyl methanethiosulfonate. To the quenched reactions, 1 volume of 2X non-reducing gel-loading solution (150 mM Tris-Cl, pH 6.8; 6% (w/v) sodium dodecyl sulfate; 25% (v/v) glycerol; 0.02% (w/v) bromophenol blue) was added. Proteins were separated by SDS-polyacrylamide gel electrophoresis and visualized by Stain-Free imaging (4-20% Criterion TGX Stain-Free, Bio-Rad #5678095). In this experiment, the fraction of cross-linked protein in the non-DTT-containing condition indicates disulfide completeness, while DTT's competitive potency reflects disulfide strength.

Negative-stain EM

GerA samples were size-exclusion-purified (TCEP was excluded for all protein variants) and diluted to $A_{280}$=0.01. Grids (CF400-CU, Electron Microscopy Sciences #215-412-8400) were glow-discharged (0.38 mBar, 30 mA, negative, glow 30 s, hold 10 s). Sample was applied to grids, followed by three washes with HBS and two with 0.75% (w/v) uranyl formate. Grids were imaged on a JEOL JEM-120i operated at 120 kV and ×60,000 magnification, and equipped with an AMT NanoSprint 15-MKII. Particles were picked, extracted, and 2D-classified without CTF correction in CryoSPARC.

Cryo-EM sample preparation and data collection

GerA samples were size-exclusion-purified and concentrated (100-kDa molecular weight cutoff). The final sample was prepared in size exclusion buffer (1 mM TCEP was included for samples that lacked a disulfide cross-link) with a protein concentration of $A_{280}$=10, along with 50 mM L-alanine, 50 mM D-alanine, or no additive. Grids (Quantifoil® R 1.2/1.3, 400 Mesh, Cu; Electron Microscopy Sciences Q450CR1.3) were glow-discharged in air using an easiGlow (Pelco) (0.38 mBar, 15 mA , negative, glow 30 s, hold 10 s). In a Vitrobot Mark IV (Thermo Fisher), 4 μL of sample was applied to the grid, followed by blotting (100% humidity, 4 °C, wait time=10 s, blot time=6 s, blot force=+23 over calibrated 0), plunge-freezing in liquid ethane held at −180 °C (Nanosoft), and storage in liquid nitrogen.

Movies were collected on a Titan Krios G3i microscope operated at 300 kV with a Selectris energy filter (10-eV slit width) and a Falcon 4i direct electron detector (Thermo Fisher) operated in counting mode (43 to 86 frames, $4{,}096^2$ pixels, ×165,000 magnification, pixel size 0.74 Å, total dose recorded in **Table S1**). Data were collected using EPU v.3.6. Three shots per hole were recorded, with a defocus range of −0.7 to −2.2 μm.

Cryo-EM data processing

All cryo-EM data processing was performed in CryoSPARC (v5.0.3) (*74*) unless indicated otherwise. Movies were motion-corrected (Patch Motion Correction) and CTF-fitted (Patch CTF Estimation). Micrographs with CTF fit resolution < 5 Å underwent template picking, using templates generated from a refined ab-initio volume derived from a subset of the same micrograph stack. Junk and on-carbon particle picks were removed (Micrograph Junk Detector), and the remaining picks were filtered by power and normalized cross correlation. Extracted particles (448-px box) underwent one round of 2D classification and two rounds of ab-initio reconstruction and heterogeneous refinement. Volume classes retained at this stage were approximately $C_5$-symmetric (though reconstructed in $C_1$) with FSC at the Nyquist resolution of the heterogeneous refinement (5.3 Å). Volume classes excluded at this stage included: uninterpretable junk; low-resolution depictions of GerA with gross aberrations such as anisotropic streaking or missing subunits; Nyquist-limited depictions of GerA with missing GerAC subunits; in the case of disulfide-containing constructs containing GerAA(A318C)/GerAC(S56C), a Nyquist-limited depiction of two 5GerAA:5GerAB:5GerAC complexes interacting with each other through a $C_2$-symmetric non-covalent interface involving one GerAC protomer from each complex. The $C_5$ symmetry of the GerAA N-terminal domains was visible in all interpretable volumes.

Retained particles underwent non-uniform refinement (*75*) in $C_5$ against a reference volume in a standardized position within the box, with optimization of per-particle defocus and per-group CTF parameters (21 groups corresponding to the 21 foil holes shot from each stage position), including spherical aberration and tetrafoil (these settings were used for all non-uniform refinements unless indicated otherwise). Particles then underwent re-extraction (448-px box) at their new centers, non-uniform refinement, reference-based motion correction (*76*), and non-uniform refinement, yielding a particle stack with a bimodal scale factor distribution. This stack then underwent three cycles of removal of low-scale-factor particles followed by non-uniform refinement, yielding Reconstruction A (the "consensus map"), which had a unimodal scale factor distribution.

This particle stack was symmetry-expanded in $C_5$ and subjected to 3D classification (10 classes, 3-Å filter, hard classification) with a mask around one GerAB protomer, yielding classes depicting GerAB in various states of mostly rigid-body rotation and translation. Classes were ordered by the within-mask GSFSC of equal-sized particle subsets, and a cutoff for class retention was applied based on manual map inspection. Retained volumes were lowpass-filtered to 3 Å and their within-mask maps were aligned to a single reference GerAB map using ChimeraX (v1.11.1) fitmap (*77*), largely removing the inter-class rotations and translations. The fitmap-calculated transformations for each class were applied to the corresponding particle stacks. Particles were then shifted to place GerAB at the center of the box, and local refinement was performed with a Gaussian prior (standard deviation 1° rotation, 0.3 Å shift), yielding Reconstruction B1. A similar protocol was applied to GerAC, yielding Reconstruction C1. Masked GerAB and GerAC protomers in Reconstructions B1 and C1 were then aligned to the corresponding density in Reconstruction A using ChimeraX fitmap. The fitmap-calculated transformations were applied to the corresponding particle stacks, and Reconstructions B1 and C1 were rebuilt in their new positions using Homogeneous Reconstruction Only, yielding Reconstructions B2 (the "GerAB-focused map") and C2 (the "GerAC-focused map").

<u>Cryo-EM model building, model refinement, and sharpened/composite map creation</u>
Model building and refinement were performed in ChimeraX (v1.11.1), Phenix (v2.1) (*78*), Coot (v0.9.8, v1.3.1) (*79*), and PyMOL (v3.1.8) (Schrödinger, LLC). For the structure of D-alanine-bound GerAA(A318C):GerAB:GerAC(S56C), individual protomers from an AlphaFold2 model of a 5GerAA:5GerAB:5GerAC complex were used as starting models, with residues mutated to cysteine where applicable. An unknown density straddled by GerAA transmembrane helices 2 and 3, likely a lipid tail, was modeled as tetradecane. An unidentified toroidal feature within the channel lumen, in line with $C_\alpha$ of Q354, was left unmodeled. The N-terminal cysteine modification of GerAC was modeled according to the chemical structure of a thioether-linked diacylglycerol previously depicted for *B. subtilis* lipoproteins (*80*), with stereochemistry around the chiral carbon assuming a parent

compound of *sn*-glycerol-3-phosphate. The acyl chains were represented by acetoxy groups. While acetylation of the N-terminal cysteine's α-amino group has been previously detected in some *B. subtilis* lipoproteins (*80*), the EM density favored modeling of this group as unmodified. The diacylglycerol modification was manually built in Coot LIDIA (v0.8.9) (*79*), and restraints for its linkage to the protein were built using AceDRG (*81*).

In early modeling rounds, phenix.real_space_refine (rigid-body docking, morphing, NQH flips, and minimization) was used to fit one GerAA protomer into Reconstruction A, one GerAB protomer into Reconstruction B2, and one GerAC protomer into Reconstruction C2 (the "primary" asymmetric unit). Refinement against the unsharpened maps (minimization only) was then alternated with manual model adjustment in Coot, guided by the global B-factor-sharpened maps from CryoSPARC. Secondary structure restraints were applied to regions of the model sitting in low-quality density (GerAA residues 355-359; GerAC residues 291-302, 235-246). In later rounds of refinement, dihedral and bond angle restraints were strengthened ad hoc for structural elements that would otherwise be dragged into outlier conformations that were not sufficiently supported by the density. Several structural elements could be seen populating multiple conformations. Alternative conformations of individual amino acid residues were built into the model if they could both exist in the same geometric context. If changing conformation of a single residue required changes to a network of neighboring residues, as was often the case in the Switch 2 region, only the conformation consistent with the local geometry was modeled.

The three models were combined and symmetry-expanded according to the $C_5$ symmetry operator present in Reconstruction A (phenix.map_symmetry, phenix.apply_ncs). The three maps were sharpened using the combined symmetry-expanded model (phenix.map_sharpening with shell sharpening, local_sharpen, model_sharpen). Reconstructions A and B2 were sharpened anisotropically. Reconstruction C2 was sharpened isotropically because anisotropic corrections degraded map quality. The sharpened maps were stitched together and symmetrized (phenix.combine_focused_maps), with the primary protomer from each model contributing its corresponding map symmetrically (weight=1). Water molecules were built into the resulting map (phenix.douse: cc_mask_filter=False, map_threshold_scale=0.5 to 0.7). A minimal asymmetric unit of water molecules surrounding the primary protomers was calculated (custom script), and the list was culled by manual inspection in Coot. One peak detected by phenix.douse near the ligand-binding site in GerAB was reassigned as $Na^+$ on the basis of its ranking by CheckMyMetal (v2.1) (*82*) (tie between $Na^+$ and $Ca^{2+}$) and its presence in the cryo-EM buffer. The symmetry-expanded, water/$Na^+$-including model was iteratively refined into the composite sharpened map. Rather than using non-crystallographic symmetry restraints (which do not yield perfect symmetry and often caused phenix.real_space_refine to crash), symmetry was enforced throughout refinement by symmetry-expanding the primary asymmetric unit at every refinement iteration. With the resulting model, occupancies and atomic displacement parameters were refined, and the above protocol for map sharpening was repeated. The re-sharpened maps were stitched together and symmetrized as above (phenix.combine_focused_maps), but this time with local weighting (local_residues=10) to optimize resolution at subunit interfaces. The combination weights calculated on the sharpened maps were also applied to the unsharpened maps, yielding the final composite maps presented in this manuscript.

While the GerAA V362 sidechain rotamer is well supported by the EM density, resolution at the sidechains of other pore helix residues (GerAA Q354, I356, L358, Q366, and E370) is poor, indicating conformational heterogeneity. Because these residues sit near the receptor's $C_5$ symmetry axis, arbitrary changes to rotamer modeling decisions, when symmetrized, seem to imply drastic changes to the channel pore profile, despite the lack of EM density support for specific rotamers. To avoid such an implication and to emphasize the heterogeneous nature of the pore-facing sidechains, we chose to model an asymmetric rotamer ensemble based on an arbitrary post-equilibration frame from the GerAA MD simulations. This frame's asymmetric pore helices (GerAA residues 350-372) were substituted into the symmetric cryo-EM model, and simulated annealing was performed (phenix.real_space_refine) with restraints targeting all dihedrals of the original cryo-EM model except select sidechain (GerAA Q354, I356, L358, Q366, and E370) and backbone (GerAA Q354) dihedrals. The pore helices were then excluded from re-symmetrization during all subsequent refinement iterations. Model coordinates were refined again, followed by refinement of occupancies and atomic displacement parameters.

With this approach, the final composite sharpened map is symmetric, and the final model is symmetric everywhere except the pore helices. For other cryo-EM datasets, models were built and refined analogously, using the D-alanine-bound GerAA(A318C):GerAB:GerAC(S56C) model as a starting model. In models lacking the GerAA-GerAC disulfide, the GerAC N-terminus was built into Reconstruction A.

Fourier shell correlation calculations

For each dataset, a preliminary hard mask was created for each protein subunit by generating a 15-Å representation of the atomic model (ChimeraX molmap), followed by binarization at a threshold that encompassed the van der Waals surface of every model atom. For GerAA, the mask encompassed all five protomers. For GerAB and GerAC, the mask encompassed only the primary protomer. These masks then underwent default soft-padding and auto-tightening in CryoSPARC. The reported FSC in the GerAA region was calculated by CryoSPARC on the half-maps of Reconstruction A; GerAB, Reconstruction B2; GerAC, Reconstruction C2. The reported FSC of the full complex was calculated on the half-maps of Reconstruction A using a composite mask generated by combining and symmetrizing the masks from the individual GerAA, GerAB, and GerAC calculations. All deposited FSC curves are the version corrected by high-resolution phase randomization, which closely tracked with the uncorrected curve at the 0.143 threshold. All resolution values mentioned in the text refer to the 0.143 crossing of the corrected FSC curve for the full complex unless indicated otherwise.

Sporulation efficiency measurements

Sporulation in liquid medium was induced by nutrient exhaustion in supplemented DS medium (DSM) (*83*) at 37 °C. After 24 hours, sporulation efficiency was determined as the total number of heat-resistant (80 °C, 20 min) colony-forming units (CFUs) divided by the wild-type strain's heat-resistant CFUs. All reported sporulation efficiency values are the mean of at least two biological replicates.

Phase-contrast microscopy of bacterial spores

24-hour-sporulated cultures were centrifuged (10,000 × *g*, 1 min), and the pellet was applied to 2% agarose pads. Phase-contrast microscopy was performed using an Olympus BX61 microscope equipped with an UplanF1 100× phase-contrast objective and a monochrome CoolSnapHQ digital camera (Photometrics) (0.0642 µm/pixel). Images were collected using MetaMorph software. Presented images depict one of the two biological replicates that were performed.

Spore preparation and purification from solid media

Strains were grown on DSM agar plates. After 96 h at 30 °C, spores were scraped from the plates, washed four times with $ddH_2O$, and resuspended in 350 µL 20% (w/v) Histodenz (Sigma-Aldrich D2158). The spore suspension was layered on top of 1 mL 50% (w/v) Histodenz and centrifuged (16,000 × *g*, 30 min) to separate dormant (phase-bright) spores (pellet) from vegetative cells, germinated (phase-dark) spores, and cell debris (interface). The pellet was washed five times with $ddH_2O$ and resuspended in 1 mL $ddH_2O$. Spores were stored at 4 °C and assayed for germination the following day.

Germination kinetics measurements

Purified spores were diluted in $ddH_2O$ to an $OD_{600}$ of 1.2, heat-activated at 70 °C for 30 min, and then incubated on ice for 20 min. The spore suspension was transferred to a clear, flat-bottom 96-well plate (80 µL per well), and an equal volume of L-alanine (2X final concentration), L-alanine/D-alanine mixture (2X final concentration), or $ddH_2O$ was added. $OD_{600}$ was recorded every 2 min using an Infinite M Plex plate reader (Tecan) (37 °C, 3 h, constant agitation between measurements). Presented data depict one of the two biological replicates that were performed (each in technical triplicate).

Immunoblot analysis

Purified spores, diluted to an $OD_{600}$ of 15 in 500 µL cold $ddH_2O$ with 1 mM PMSF, were transferred to tubes containing lysis matrix B (MP Biomedicals #116911050) and cooled on ice. Spores were lysed using a FastPrep (MP Biomedicals) (6.5 m/s, 60 s), followed by addition of 500 µL 2X sample buffer (4% (w/v) SDS; 250 mM Tris, pH 6.8; 20% (v/v) glycerol; 10 mM EDTA; 0.02% (w/v) bromophenol blue; 10% (v/v) β-mercaptoethanol). After centrifugation (20k × *g*, 5 m), supernatant was collected. An equal volume of each supernatant was resolved by SDS-PAGE (4-20% Mini-PROTEAN TGX, Bio-Rad #4561096) and transferred onto 0.2-µm nitrocellulose using the Trans-Blot Turbo system (Bio-Rad). Membranes were blocked in Tris-buffered saline (TBS: 50 mM Tris-Cl, pH 7.5; 150 mM NaCl) with 0.5% (v/v) Tween-20 and 5% (w/v) non-fat milk, then probed with α-GerAA (1:5,000) (*84*) or α-SpoVAD (1:10,000) (*85*) in TBS with 0.05% (v/v) Tween-20 and 3% (w/v) bovine serum albumin. Primary antibodies were detected using horseradish-peroxidase-conjugated goat α-rabbit IgG (1:4,000) (Bio-Rad #1706515) and Clarity Western ECL substrate (Bio-Rad #1705060). Presented data depict one of two biological replicates that were performed.

Molecular dynamics simulations

To build the simulation system for the GerAA pentamer's transmembrane domain, we started from a preliminary symmetric cryo-EM model of D-alanine-bound GerAA(A318C):GerAB:GerAC(S56C), with engineered cysteines reverted to their wild-type identities. Except for the GerAA-L417 rotamer, there were no ligand-dependent changes to GerAA in the cryo-EM maps, so choosing a different starting model would not be expected to meaningfully change simulation results. GerAB and GerAC subunits, as well as residues 1-272 and 433-482 of GerAA, were removed using Prime (Schrödinger) (*86*). The GerAA pentamer's pore was aligned to the *z*-axis using the Orientations of Proteins in Membranes webserver (*87*). Prime was used to add capping groups to each protein chain's termini. Protonation states of all titratable residues were assigned at pH 7. Histidine residues were modelled as neutral, with a hydrogen atom bound to either the delta or epsilon nitrogen depending on which tautomeric state optimized the local hydrogen-bonding network. Using Dabble (*88*), the prepared protein structures were inserted into a pre-equilibrated palmitoyl-oleoyl-phosphatidylcholine (POPC) bilayer, the system was solvated, and sodium and chloride ions were added to neutralize the system and to obtain a final concentration of 150 mM. The final system comprised approximately 93,000 atoms, and system dimensions were approximately 120x120x70 Å.

We used the CHARMM36m force field for proteins, the CHARMM36 force field for lipids and ions, and the TIP3P model for waters (*89–91*). All simulations were performed using the Compute Unified Device Architecture (CUDA) version of particle-mesh Ewald molecular dynamics (PMEMD) in AMBER20 on graphics processing units (GPUs).

The system first underwent three rounds of minimization, each consisting of 500 cycles of steepest descent followed by 500 cycles of conjugate gradient optimization. 10.0 and 5.0 $kcal \cdot mol^{-1} \cdot Å^{-2}$ harmonic restraints were applied to the protein and lipids for the first and second rounds of minimization, respectively. 1 $kcal \cdot mol^{-1} \cdot Å^{-2}$ harmonic restraints were applied to the protein for the third round of minimization.

We performed twelve independent simulations, with initial atom velocities assigned randomly and independently during system heating. Systems were heated from 0 K to 100 K in the NVT ensemble over 12.5 ps and then from 100 K to 310 K in the NPT ensemble over 125 ps, using 10.0 $kcal \cdot mol^{-1} \cdot Å^{-2}$ harmonic restraints applied to protein heavy atoms. Subsequently, systems were equilibrated at 310 K and 1 bar in the NPT ensemble, with harmonic restraints on the protein non-hydrogen atoms that started at 5.0 $kcal \cdot mol^{-1} \cdot Å^{-2}$ and tapered stepwise (decrease by 1.0 $kcal \cdot mol^{-1} \cdot Å^{-2}$ every 2 ns for 10 ns, then decrease by 0.1 $kcal \cdot mol^{-1} \cdot Å^{-2}$ every 2 ns for 20 ns). Production simulations were performed without restraints at 310 K and 1 bar in the NPT ensemble using the Langevin thermostat, the Monte Carlo barostat, and a timestep of 4.0 fs with hydrogen mass repartitioning (*92*). Bond lengths were constrained using the SHAKE algorithm (*93*). Non-bonded interactions were cut off at 9.0 Å, and long-range electrostatic interactions were calculated using the particle-mesh Ewald (PME) method (Ewald coefficient ~0.31 $Å^{-1}$, $4^{th}$-order B-spline interpolation). The PME grid size was chosen such that the width of a grid cell was approximately 1 Å. Frames from production trajectories were saved every 200 ps. The AmberTools17 CPPTRAJ package was used to reimage trajectories (*94*). Simulations were visualized and analyzed using Visual Molecular Dynamics (*95*) and PyMOL (v3.1.8) (Schrödinger, LLC).

Pore size calculations

MOLE (*66*) (default settings, “Radius” output) was used to calculate the diameter of the maximally inscribed ball limited by the van der Waals surface of the surrounding protein atoms.

For molecular dynamics simulation plots, a “distance” parameter was designed as a computationally inexpensive approximation of the MOLE-calculated diameter. “Distance” was calculated as the minimal pairwise distance between any two atoms (including hydrogens) of the residue of interest in non-adjacent protomers. That is, for each atom of residue *i* in protomer *n*, we calculated the distance to each atom of residue *i* in protomers *n*+2 and *n*+3, and we reported the minimum distance encountered after all atom pairs in all protomer pairs had been considered. Differences from the MOLE-calculated diameter are expected to be minor: the line segment traversing this distance is usually an off-center chord of the circle about the pore axis (decreasing the distance as compared to MOLE), but the atomic-center-based measurement does not factor in the atoms’ finite van der Waals radii (increasing the distance as compared to MOLE).

The AlphaFold predictions and the cryo-EM structures all underwent hydrogen addition and AMBER relaxation before being subjected to the same “distance” calculation, whose results are reported in **Table S2**.

### SUPPLEMENTARY NOTES

Supp. Note 1: Interpreting sporulation and germination assays

In the standard evaluation of a GerA mutant's phenotype in spores, four assays were performed on each strain. Phase-contrast light microscopy and colony-counting-based quantification of heat-resistant spores were performed on unpurified sporulated cultures (**Fig. S5**). Immunoblot-based measurements of GerAA levels (**Fig. S6**) and OD-based measurements of germination kinetics (**Fig. S5**) were performed on purified phase-bright spores. To assign a GerA mutant as either impaired or hyperactivated, all assays must be considered together, with support for assignment coming from specific combinations of results that are outlined in the table below (which uses WT GerA as a baseline for comparison).

| | **phase-contrast micrographs** | **heat-resistant spore counts ("sporulation efficiency")** | **immunoblot** | **germination kinetics (OD)** |
|---|---|---|---|---|
| **impaired GerA activity** | fewer phase-dark spores* | fewer heat-resistant CFUs | unchanged GerAA levels** | slower $v_{max}$, smaller $A$ |
| **GerA hyperactivity** | more phase-dark spores | fewer heat-resistant CFUs | unchanged or lower GerAA levels | similar $v_{max}$, shorter $t_{lag}$, similar or smaller $A$ |

*The fraction of phase-dark spores in strains containing WT GerA is already so close to zero that, while an impaired GerA mutant is expected to push the fraction even closer to zero (*96*), it is not a robust way to assess GerA activity and is mentioned here only for completeness.

**This table only considers mutants with no change to the GerAA levels for assignment of impaired GerA activity. Mutants with lower measured GerAA levels are only considered for assignments of GerA hyperactivity or uninterpretable activity. See text below.

In phase-contrast micrographs, dormant spores are phase-bright, while germinated spores are phase-dark. Appearance of phase-dark spores during sporulation is attributed to premature germination, which can result from GerA hyperactivity (*15*, *96*).

Counting post-heat-treatment CFUs ("heat-resistant spore counts" or "sporulation efficiency") measures the ability of spores to both sporulate and germinate. Impaired GerA mutants successfully sporulate and survive heat treatment but fail to germinate, yielding fewer heat-resistant CFUs. Hyperactive GerA mutants cause spores to prematurely germinate or decay into non-heat-resistant forms, also yielding fewer heat-resistant CFUs.

Because GerAA, GerAB, and GerAC depend on each other for stability in the developing spore, a decreased level of any individual subunit is expected to cause a decrease in the GerAA level measured by immunoblot (*10*, *97*, *98*). Thus, blotting for GerAA alone measures total GerA complex levels in the spore. We performed immunoblots primarily to determine whether mutations that alter germination phenotypes do so by changing the function of intact GerA complex or by destabilizing the complex. GerA-destabilizing mutations, for which impaired vs. hyperactive assignments cannot be made, are not considered in the table above. However, a decrease in measured GerAA levels does not necessarily emerge from a change to GerA complex stability in the bulk sporulated culture. For example, we observed that hyperactive GerA mutants often had lower GerAA levels, which we attribute not to GerA complex instability but to a selection effect. In populations of spores with a heterogeneous distribution of germinant receptor copy number (*99*, *100*), spores containing higher levels of germinant receptors are more likely to prematurely germinate. When phase-bright spores are purified away from prematurely germinated spores before the immunoblot, a population with lower levels of germinant receptors is selected (even before purification, prematurely germinated spores are expected to degrade GerA as they attempt to transition back to vegetative growth). When a hyperactive GerA mutant is present, the cutoff in GerA copy number for premature germination is pushed even lower, yielding a purified phase-bright spore population with lower GerAA levels. Therefore, mutants that exhibit signatures of GerA hyperactivity and a weaker α-GerAA immunoblot band can be interpreted as hyperactive *in spite of* lower copy number, and the lower copy number can optionally be interpreted as a direct result of the hyperactivity.

A simple model for OD-based germination kinetics is presented below. For impaired GerA mutants, $v_{max}$ is slower and the amplitude is smaller. Small $v_{max}$ slopes make $t_{lag}$ a noisy and unreliable parameter when assessing impaired mutants. When $v_{max}$ is similar to WT, a reduced $t_{lag}$ is the signature of a hyperactive GerA mutant. Counterintuitively, hyperactive GerA mutants often yield decreased amplitude, likely for a reason analogous to that described above for the immunoblot results. Among the heterogeneous spore types generated in any sporulated culture, phase-bright spore purification enriches the defective population that can only germinate slowly or not at all. For this reason, $t_{lag}$ is a more informative parameter for assessing hyperactivity than $v_{max}$, whose ensemble behavior is expected to emerge from per-spore contributions that are (roughly) scaled by the amplitude.

Some departures from the summary table above may be identified in the data. For example, a "hyperactive" GerA mutant that yields faster germination in the OD assay may or may not change the phase-contrast micrographs or the heat-resistant spore counts, depending on whether the hyperactivity is constitutive, heat-activated, or ligand-activated. Careful evaluation of GerA mutant behavior must consider the results of all assays together.

<u>*Modeling the drop in optical density associated with germination of a spore suspension*</u>

An example set of OD curves is shown below for reference. To provide a language for interpreting these curves, we present a simple model with descriptive parameters. Readers can quickly assess these parameters from the data by eye, but we calculate and plot them rigorously in this example for clarity. For each curve, we performed least-squares line fitting to every 24-minute sliding window and selected the line with the greatest slope (this slope is $v_{max}$). The time at which this line intersects the line y=1 is labeled $t_{lag}$. The difference between 1 and the OD value at t=3 h (which is independent of the line-fitting process) is labeled *A* (amplitude). The true plateau value would be a more informative parameter than this arbitrarily defined "amplitude" but is usually unconstrained by the data collected within the 3-h window. A more sophisticated model, involving convolution of a Gamma-distributed $t_{lag}$ with two exponential decay terms, is required to fit the full sigmoid, but it provides no interpretive benefit over the simpler linear model.

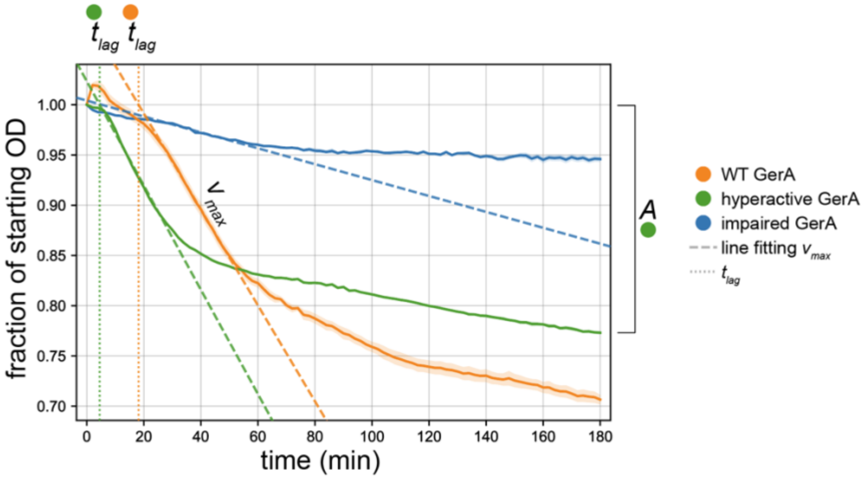

Supp. Note 2: Explaining the different local structural consequences of L- vs. D-alanine binding by GerAB

It was not immediately apparent why TM6A responds so differently to L- and D-alanine (**Fig. 2A**), given each isomer's amine contacts the same TM6A backbone carbonyls (**Fig. S8**). One possible cause of the difference is GerAB-L199, whose sidechain rotamer must flip to accommodate the sidechain methyl of L- but not D-alanine (**Fig. S8**). Interestingly, the L-alanine-specific rotamer is compatible with the adjacent TM6A helix only in its unraveled form. However, when we removed L199's potential to cause rotamer-specific effects by mutation to alanine, it resulted in an increase in phase-dark spores and only minor changes to sporulation efficiency and germination kinetics (**Fig. S5**). The L199G mutation yielded similar results but with a more intense attenuation of sporulation efficiency (**Fig. S5**). These results argue that the L199 rotamer flip does not cause TM6A unraveling and likely plays a more nuanced role.

Instead, we note that the shared carboxylate, $C_\alpha$, and amine of L-alanine and D-alanine occupy slightly different poses within GerAB, tilting in the direction expected from collision of the sidechain methyl with the walls of the sidechain-sensing cavity (**Fig. S8**). Thus, L-alanine tilts further toward GerAA than D-alanine does. The result is that L-alanine pulls the base of TM6A closer to GerAA, inducing a greater distortion that can likely only be relieved by helix unraveling (**Fig. 2A**). Tilting of L-alanine toward GerAA is part of the same motion that activates Switch 2, which involves movement of several ligand-proximal structural features toward GerAA (**Fig. S10A**). Interestingly, 2-aminoisobutyrate, which combines the sidechain methyls of L- and D-alanine, is also a germination initiator (presumably acting through GerA) with similar efficacy to L-alanine but 15- to 20-fold worse potency (*11*, *12*). Glycine, which has neither sidechain methyl, is a competitive germination inhibitor with 10- to 20-fold worse potency than D-alanine (*11*, *12*). We propose that the efficacy of each ligand emerges from its ability to access TM6A-distorting tilt angles, while potency emerges from the coupling of these tilt angles to the energetic landscape of protein conformational changes, as well as the energy of the contacts between the protein and the ligand sidechain methyls.

Supp. Note 3: Additional structural features controlling Switch 2 propagation
Here we expand on the analysis of structural features that control Switch 2 propagation. However, for clarity of the following discussion, we first mention a difference between the L-alanine-bound GerAA(A318C)-GerAC(P52C) structure and the L-alanine-bound GerAA(A318C)-GerAC(S56C) structure in the region where GerAB contacts the GerAA transducer helix. Specifically, the different disulfides cause two GerAC loops (residues 88-97, 119-126) to pack in different ways (**Fig. S9A**), which results in minor changes to some GerAB sidechains in this region. As neither of these GerAC loops contains the disulfide, we posit that a higher-order feature of the structure, such as GerAC's seesaw motion, causes the repacking. The alternative loop conformations may underlie GerAA(A318C)-GerAC(P52C)'s greater occupancy of Switch 2's activated state as compared to GerAA(A318C)-GerAC(S56C) (**Fig. 3D**), at least in the detergent-solubilized context.

We begin our supplementary discussion of Switch 2 activation in the middle of the rotated GerAB-TM8 segment. In apo GerA, the GerAB-L276 sidechain fits into a pocket formed by GerAB-L160, T255, I256, and F259. Upon L-alanine addition, GerAB-L276 pulls out of the pocket into an unrestricted position with no van der Waals contacts (**Fig. S9B**). Mutating GerAB-L276 to alanine resulted in impaired germination (**Fig. S5**), perhaps by enabling the sidechain at position 276 to nestle more snugly into the pocket in TM8's inactive conformation. GerAB-L277 is held in a similar hydrophobic pocket (**Fig. S9C**), but its mutation caused a decrease in GerAA levels (**Fig. S6**) without an obvious hyperactivity phenotype (**Fig. S5**), making its effect on GerA signaling uninterpretable (**Supp. Note 1**).

We next discuss a GerAB/GerAC contact that explains GerAC's ability to allosterically enhance GerAB-TM8 rotation: a salt bridge between GerAC-R125 and GerAB-E273, which sits at the top of GerAB-TM8's rotated segment (**Fig. 3B**, **Fig. S9D**). This interaction is important for GerA complex stability, as mutation of either residue to alanine caused loss of GerAA in spores (**Fig. S6**, **Supp. Note 1**). However, the more conservative GerAB(E273D) mutation maintained GerA complex stability (**Fig. S6**) but strongly attenuated germination (**Fig. S5**), highlighting this interaction's importance for Switch 2 activation in addition to complex stability. GerAC-R125 is braced by a hydrogen-bonding network involving GerAB-E263 and GerAC-S127 (**Fig. S9D**). GerAB(E263A) impaired GerA activity, while GerAC(S127A) hyperactivated (**Fig. S5**), arguing for a role of this network in regulating Switch 2 activation.

In our model, the consequence of GerAC-enhanced GerAB-TM8 rotation is a movement of the adjacent GerAA transducer helix (residues 310-320). This signal transmission could occur, at least partially, via the contact between GerAB-S274 and GerAA-L307, which lies in the loop at the base of the transducer helix (**Fig. S9E**). The GerAB(S274G) and GerAA(L307A) mutations each reduced sporulation efficiency by 60-70%, supporting the importance of this contact in signal transmission (**Fig. S5**).

Finally, we consider the extracellular loop extension of GerAB-TM8 (residues 263-271), which we term here the TM8 hairpin loop (**Fig. S9E**). The hairpin loop was previously shown to be a hotspot for mutational suppressors of a GerAA hypermorph (*15*), and the residue that was mutated in the hypermorph (GerAA(P326S)) directly contacts the hairpin loop in our structures (**Fig. S9F**). We explain the importance of the hairpin loop in terms of its numerous contacts with the base of the GerAA transducer helix (**Fig. S9F**). The backbone of the hairpin loop makes van der Waals contacts to GerAA-P309. Additionally, GerAB-R271 forms GerAA(A318C)-GerAC(P52C)-specific contacts with the carbonyl oxygen of GerAA-L308 and the adjacent GerAB residue E270, which itself points toward the positive end of the transducer helix dipole. In this conformation, GerAB-R271 also contacts the above-mentioned GerAB-E263 residue involved in GerAC-R125 positioning (**Fig. S9F**). As mutation of GerAB-R271 to alanine reduces GerA complex levels in spores (**Fig. S6**), we cannot draw firm conclusions about its role in Switch 2 activation, but we suggest that these contacts transfer the force generated by TM8 rotation into the base of the GerAA transducer helix. On the distal side of the TM8 hairpin loop, GerAB-L264 tucks into a hydrophobic pocket formed by GerAB-K169, F262, and F272 (**Fig. S9F**). GerAB(L264A) resulted in modestly hyperactive GerA phenotypes (**Fig. S5**), suggesting that this sidechain's residence in the pocket holds the hairpin loop in an inactive conformation.

Interestingly, the GerAB-TM8 hairpin loop is disordered in WT GerA but ordered in the disulfide-stabilized GerA complexes (**Fig. S9G**), in a ligand-independent manner. Thus, in addition to allosterically enhancing GerAB-TM8 rotation, GerAC may also function to stabilize the TM8 hairpin loop, perhaps enabling further propagation of Switch 2 in a fully activated complex.

Together, these structural observations provide hints as to how nutrient-induced TM8 rotation causes movement of the transducer helix, which induces channel opening at the complex center.

Supp. Note 4: GerAC toe-heel docking

To facilitate discussion of GerAC conformation, we point out that the global structure of a single GerAC protomer resembles a foot (**Fig. S13**). A key difference between the cryo-EM structures and the AlphaFold predictions involves an interaction between the "toe" of one GerAC protomer and the "heel" of the adjacent protomer. In the "undocked" state of the cryo-EM structures, the toe is separated from the adjacent heel, and several non-contiguous stretches of amino acid residues in the toe are disordered. In the "docked" state, the toe packs directly against the heel of the adjacent protomer. This interaction is accomplished by global rotation of each GerAC protomer, along with extension of the toe's β-sheet to several residues past where the sheet ends in the cryo-EM structures (**Fig. S13**).

In the docked state, the GerAC toe interacts not only with the adjacent GerAC heel but also with the top of GerAB (**Fig. S13**). To test the possibility that toe-heel docking occurs *in vivo*, we mutated two pairs of residues that interact only in the docked state.

For one pair (GerAB-E119/GerAC-R275), which forms a salt bridge in the docked state (**Fig. S13**), we performed a charge swap. GerAB(E119R) behaved similarly to WT, GerAC(R275E) exhibited modest GerA hyperactivity, and combining the mutations yielded even stronger GerA hyperactivity (**Fig. S5**), despite extensive depletion of GerAA levels in the purified spores (**Fig. S6**, **Supp. Note 1**). These results do not fit the neat behavior of a classic salt bridge interaction that was demonstrated for GerAB-E182/GerAC-K339 (**Fig. 2D**). However, they do suggest that GerAB-E119 and GerAC-R275, either separately or together, play an important role in GerA function that would not be predicted from their solvent-exposed positions in the undocked state.

The second residue pair (GerAB-R33/GerAC-F287) forms a cation-π interaction in the docked state (**Fig. S13**). We expected that the GerAC(F287D) mutation would hold GerAB-R33 more stably, while the GerAC(F287R) mutation would destroy the interaction, perhaps yielding opposite functional results. Instead, both mutations yielded similar levels of GerA hyperactivation (**Fig. S5**), suggesting that the GerAB-R33/GerAC-F287 interaction holds GerA in an inactive state *in vivo* and that any geometric disruption causes activation. Independent of the precise mechanistic interpretation, the fact that the GerAC-F287 mutations have such strong phenotypes supports the idea that, *in vivo*, GerAC populates conformations besides that of the undocked cryo-EM structures, in which F287 is completely disordered and has no obvious function (**Fig. S13**).

These observations argue that GerAC toe-heel docking could be important for GerA function. Currently, it is unclear whether the GerAC docking (or undocking) transition is part of *in vivo* GerA activation or if the docked state is a constantly-occupied prerequisite of full channel opening. If the latter, the undocked state could be an off-pathway *in vitro* artifact that would explain the partially activated conformation occupied by the cryo-EM structures.

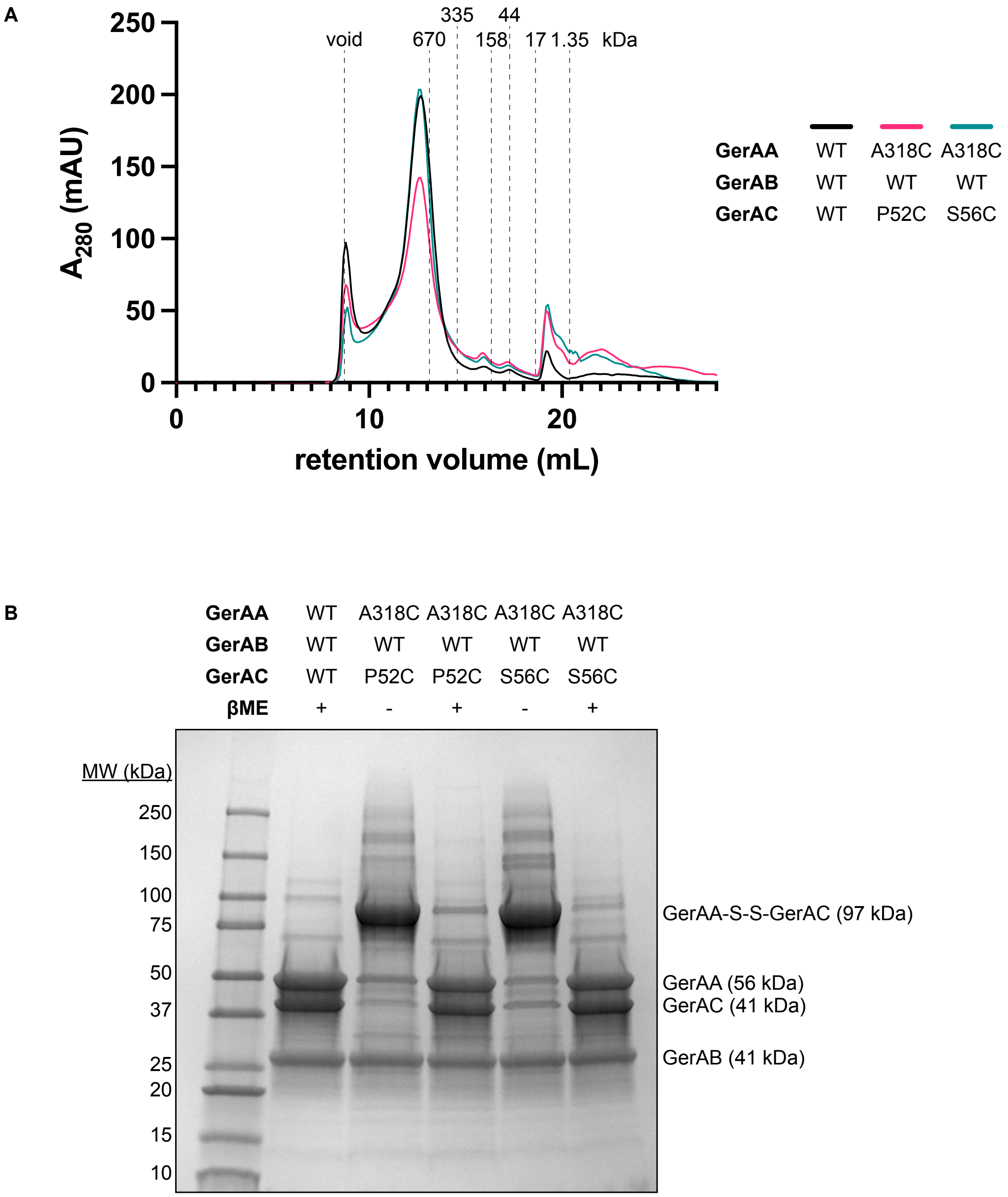


**Fig. S1.** (**A**) UV absorbance trace of size exclusion chromatography of cryo-EM samples. (**B**) One-Step-Blue-stained SDS-PAGE analysis of cryo-EM samples. As expected, integral membrane proteins GerAA and GerAB have greater electrophoretic mobility than soluble protein standards of comparable molecular mass.

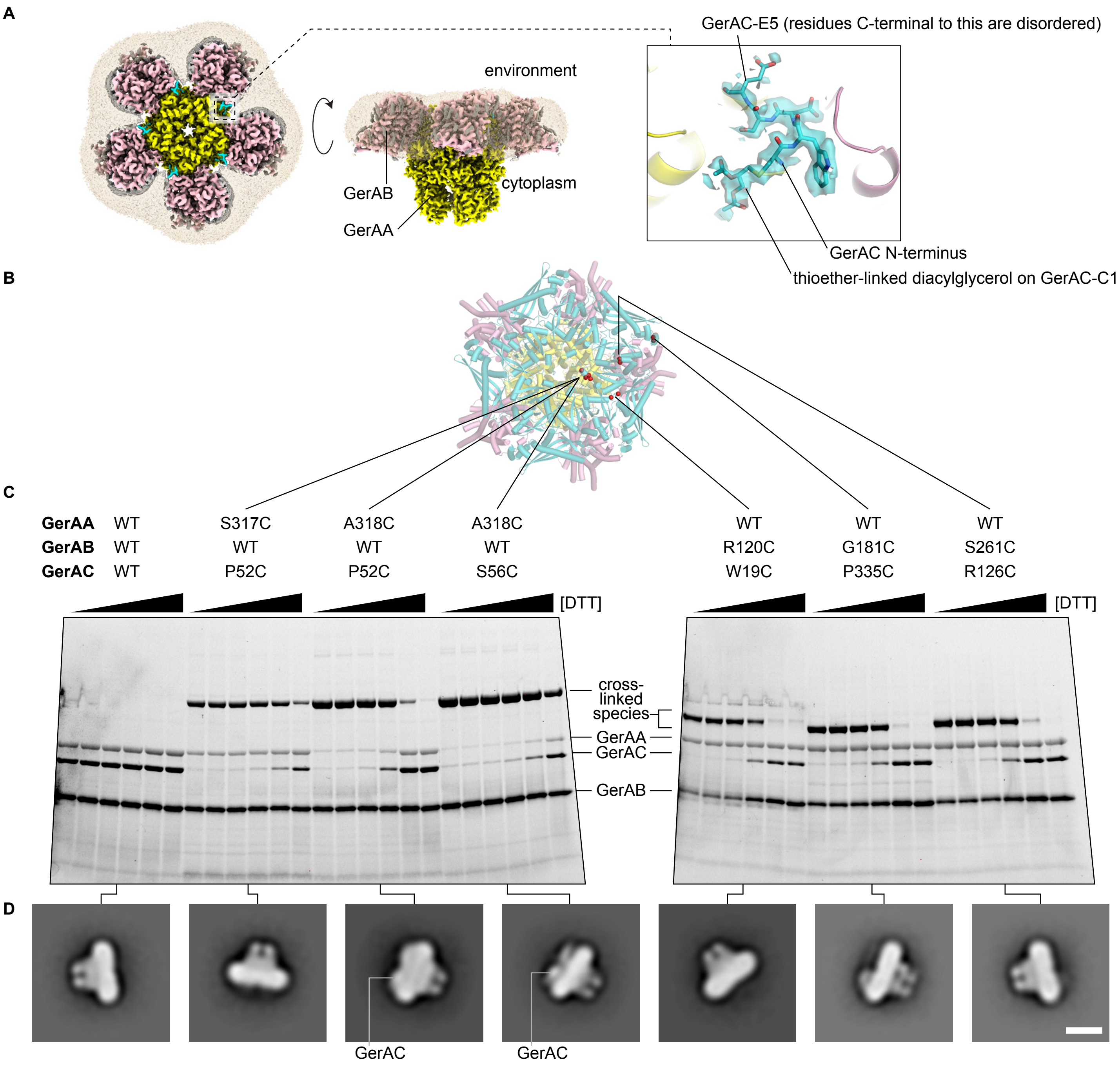


**Fig. S2.** (**A**) WT GerAA:GerAB:GerAC, apo, unsharpened composite map. Ordered detergent, opaque gray. Disordered detergent, translucent beige (unsharpened consensus map). Inset, detail of five ordered N-terminal residues of GerAC, with lipid modification (sharpened consensus map). (**B**) AlphaFold-predicted model of 5GerAA:5GerAB:5GerAC with locations of engineered disulfides highlighted as red spheres. **(C)** SDS-PAGE/Stain-Free analysis of purified GerA complexes exposed to DTT (0, 0.01, 0.1, 1, 10, 100 mM). **(D)** Negative-stain EM 2D class averages depicting side view of purified GerA complexes. Scale bar, 10 nm.

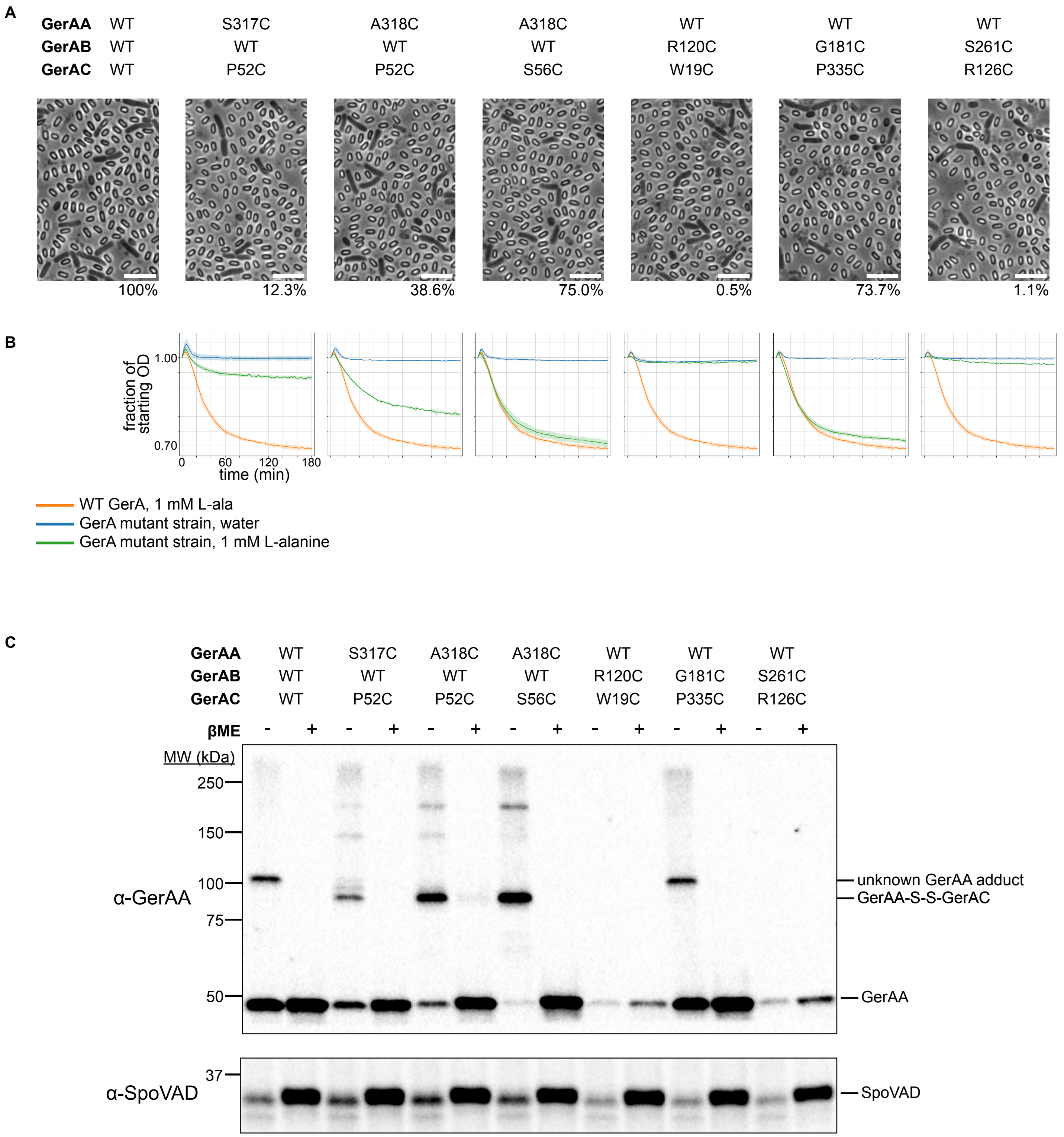


**Fig. S3.** (**A**) Phase-contrast light micrographs of sporulated cultures expressing GerA mutants with engineered cystine cross-links (see **Fig. S2**). The percentage underneath is the average sporulation efficiency, defined in the Methods and discussed in **Supp. Note 1**. (**B**) Spore germination kinetics in response to L-alanine addition, as measured by OD drop (strains indicated in (A)). (**C**) Immunoblots assessing GerAA levels and extent of cross-linking in the spores from (A). SpoVAD, loading control.

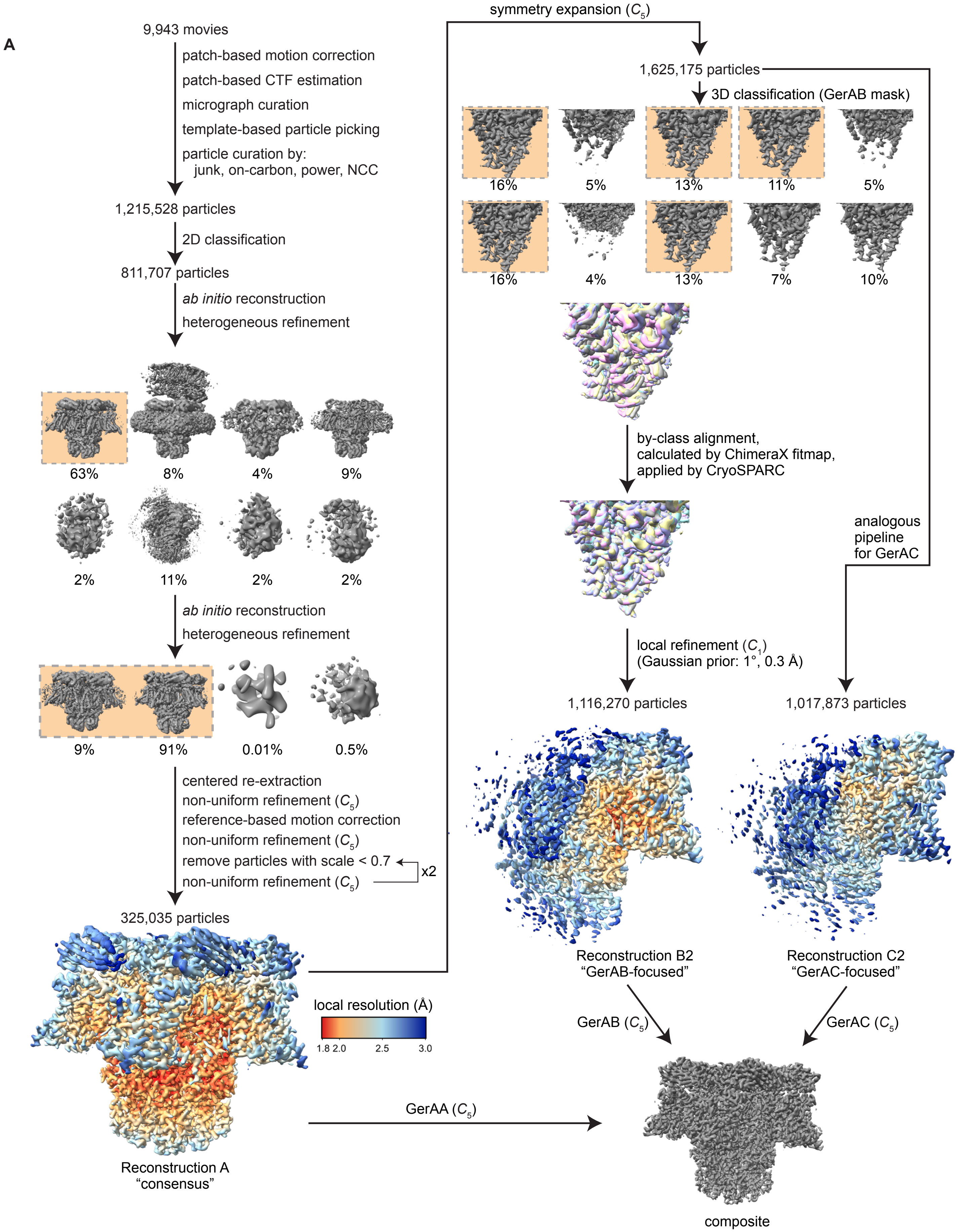


**Fig. S4 (page 1/2).** (**A**) Cryo-EM data processing scheme for GerAA(A318C):GerAB:GerAC(S56C) bound to D-alanine. Other cryo-EM datasets were processed analogously. The "8%" class in the first round of heterogeneous refinement is the $C_2$-symmetric dimer of pentamers described in the Methods. Only unsharpened volumes are depicted in this figure.

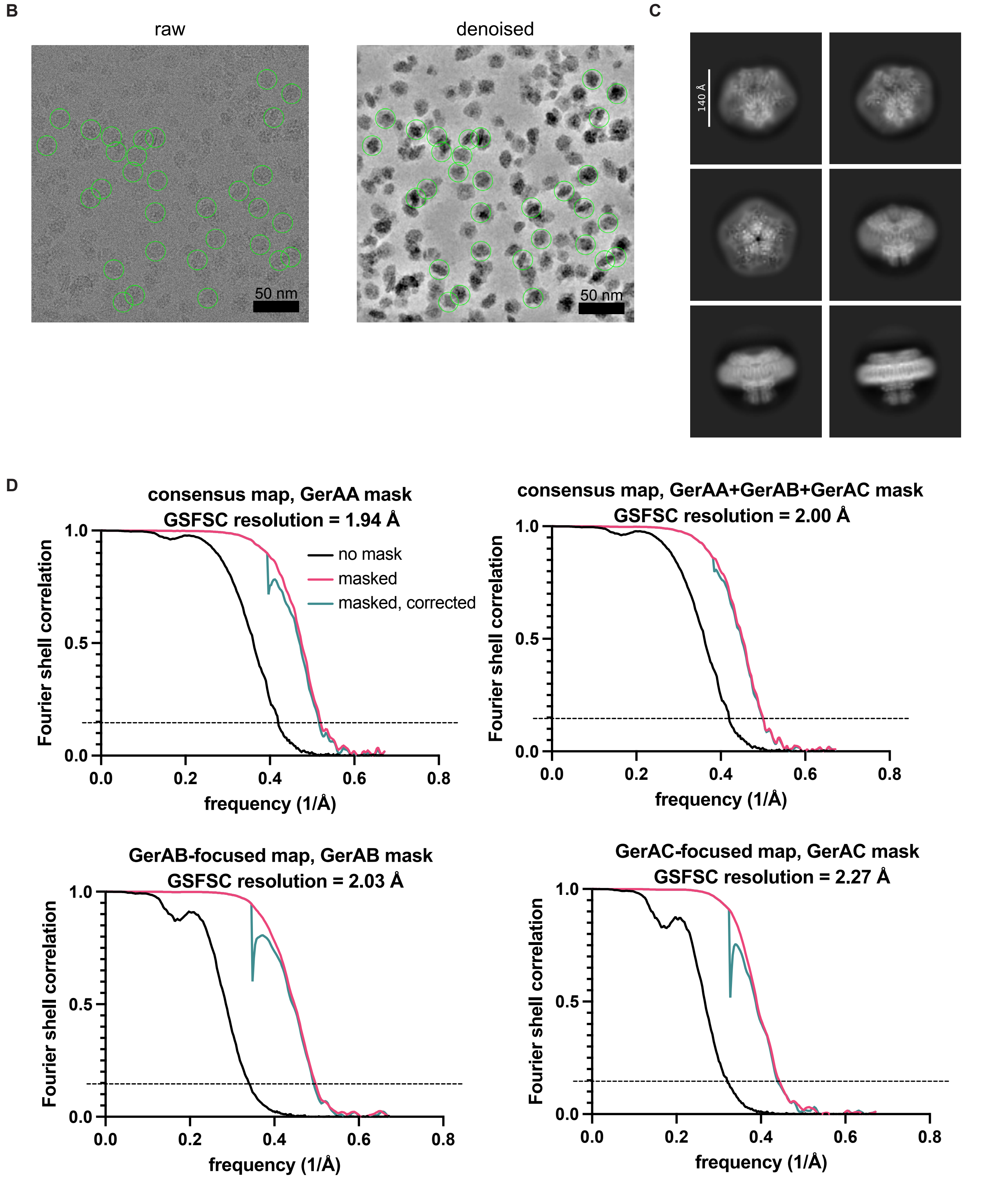


**Fig. S4 (page 2/2).** (**B**) Left: example motion-corrected micrograph. Right: the same image subjected to CryoSPARC's Micrograph Denoiser. Particles present in the final consensus stack are circled in green. (**C**) Example 2D class averages from the particle curation stage. (**D**) Gold-standard Fourier shell correlation (GSFSC) curves with correction by phase randomization. The horizontal line represents the FSC = 0.143 threshold. FSC data for other cryo-EM datasets are available in **Table S1** and the EMDB depositions.

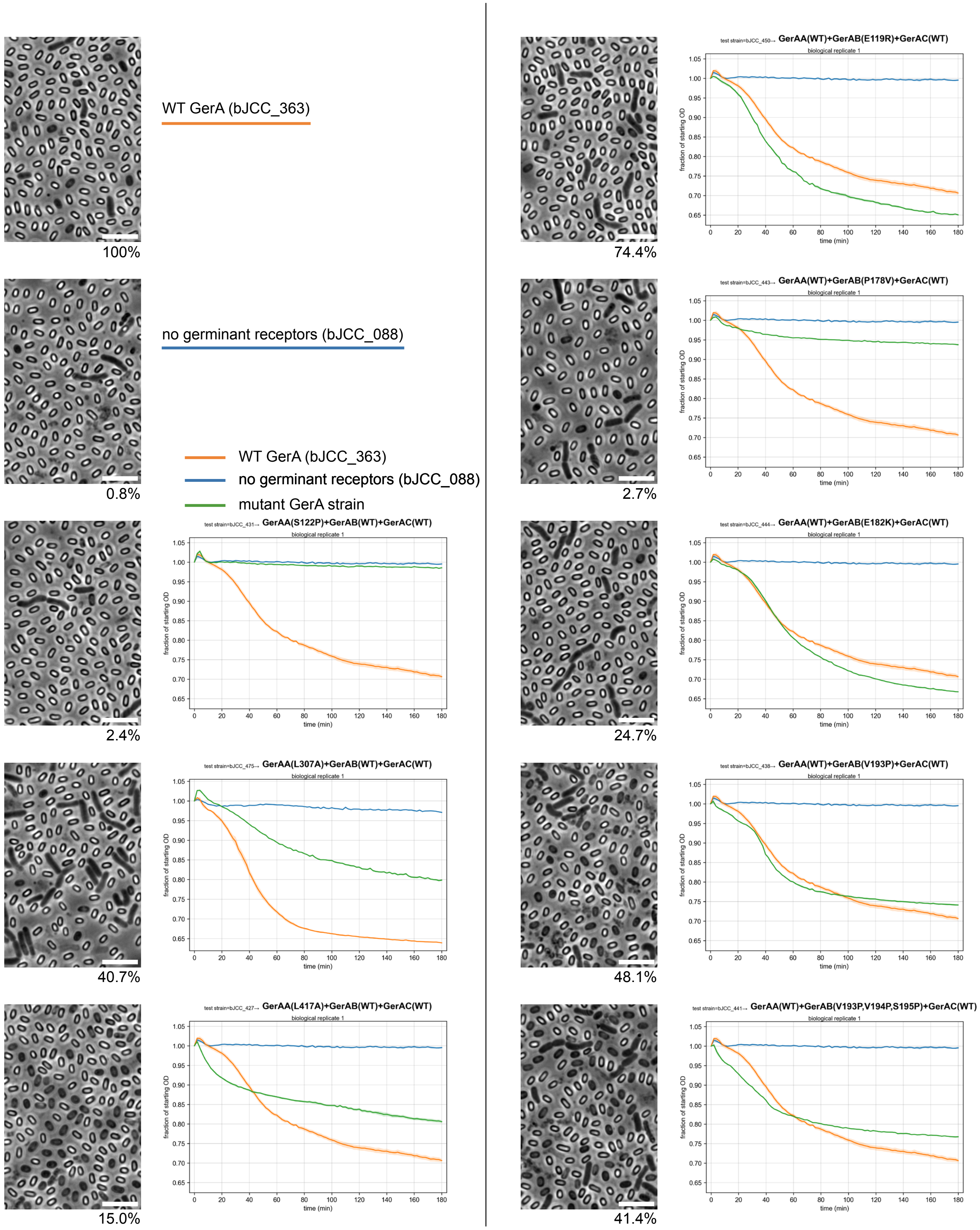


**Fig. S5 (page 1/4).** Left, phase-contrast light micrographs of sporulated cultures expressing WT GerA, no germinant receptors, or GerA mutants. Scale bars, 5 µm. The percentage underneath is the average sporulation efficiency, defined in the Methods. Right, germination kinetics of spores purified from the culture depicted in the micrograph. Shaded area, standard error of the mean for three technical replicates. See **Supp. Note 1** for details of assay interpretation. Strains are presented in order of mutated subunit (GerAA, GerAB, GerAC) then residue number. Strains with mutations in two subunits are presented at the end.

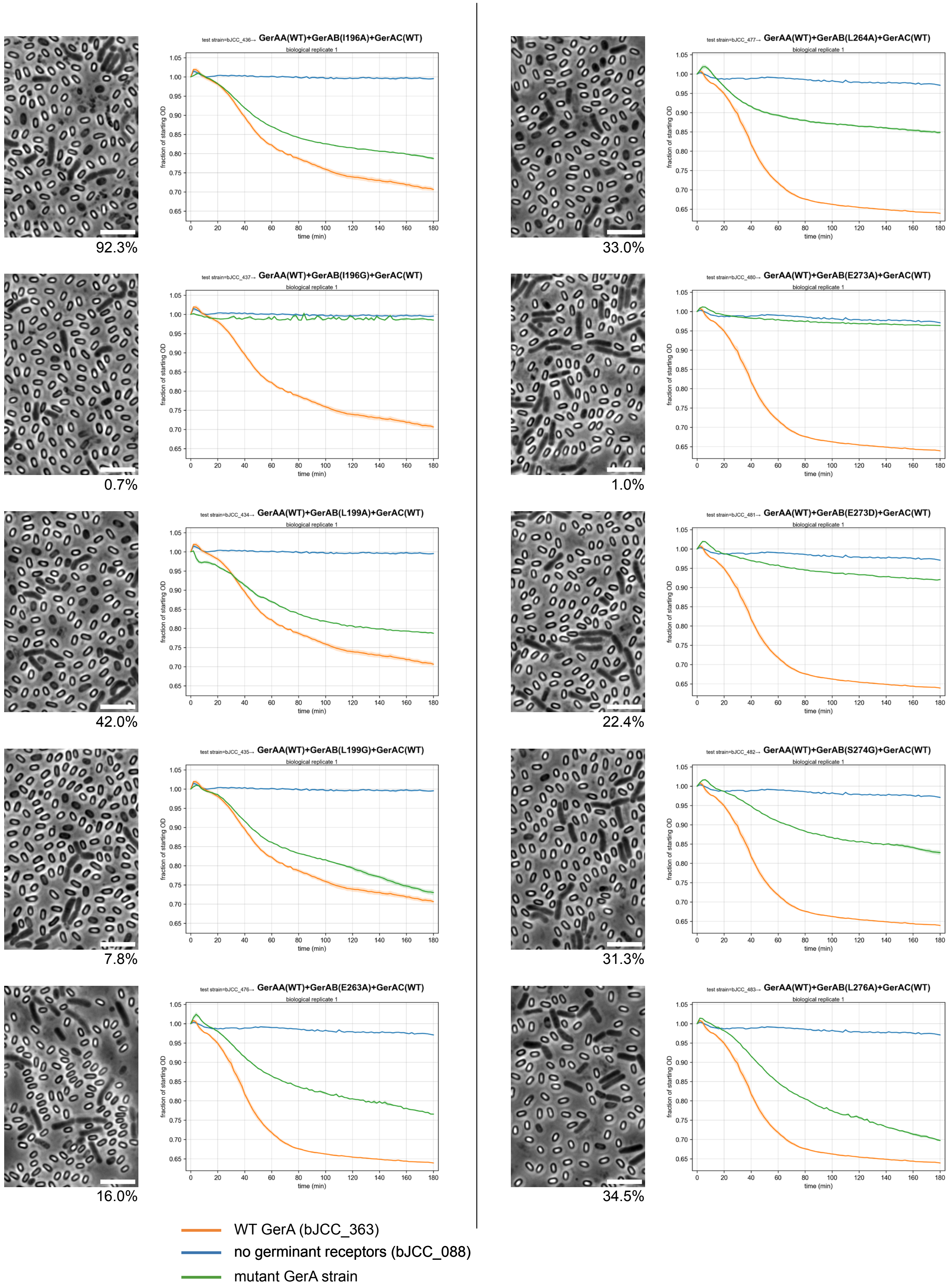


**Fig. S5 (page 2/4).**

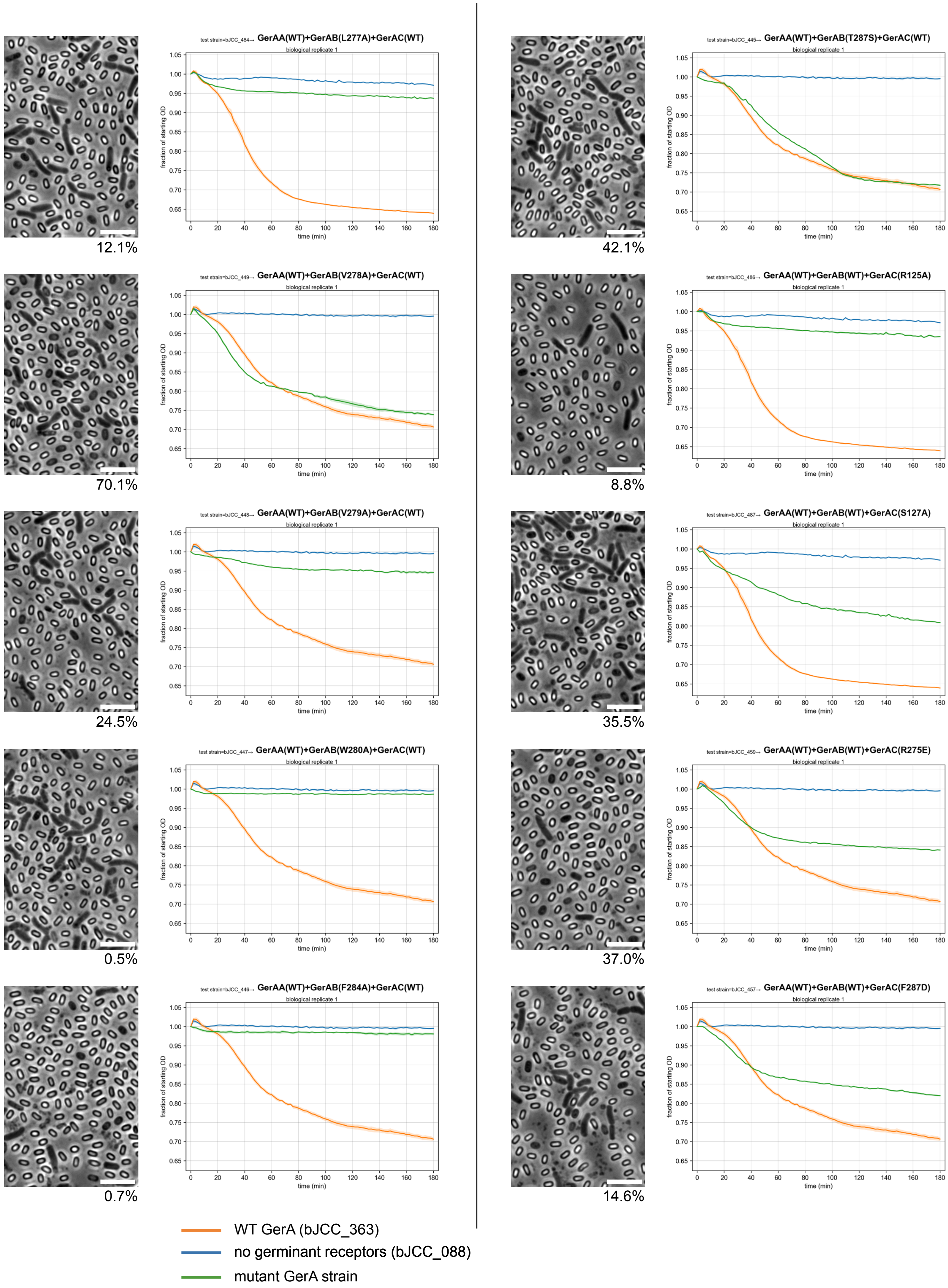


**Fig. S5 (page 3/4).**

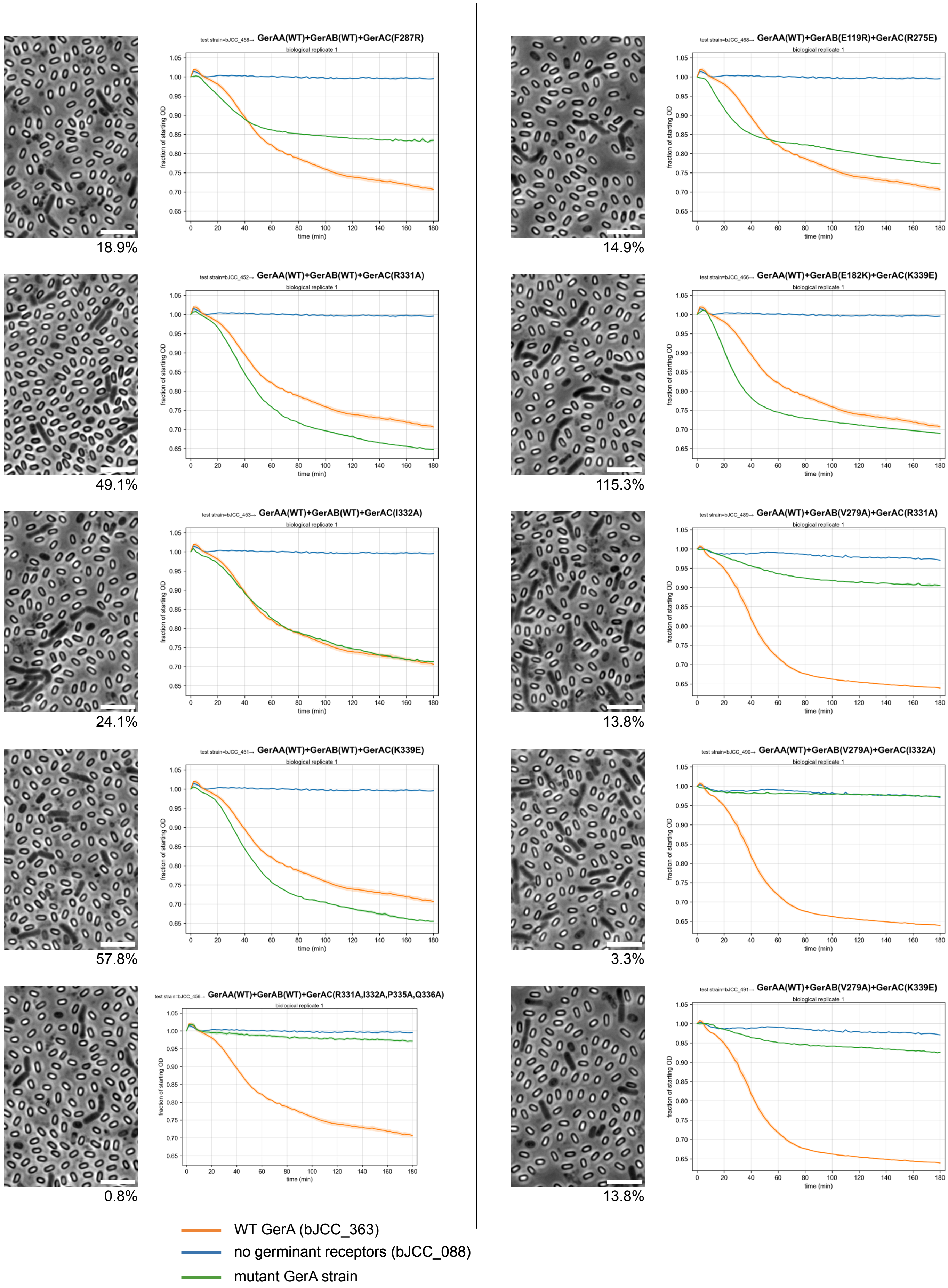


**Fig. S5 (page 4/4).**

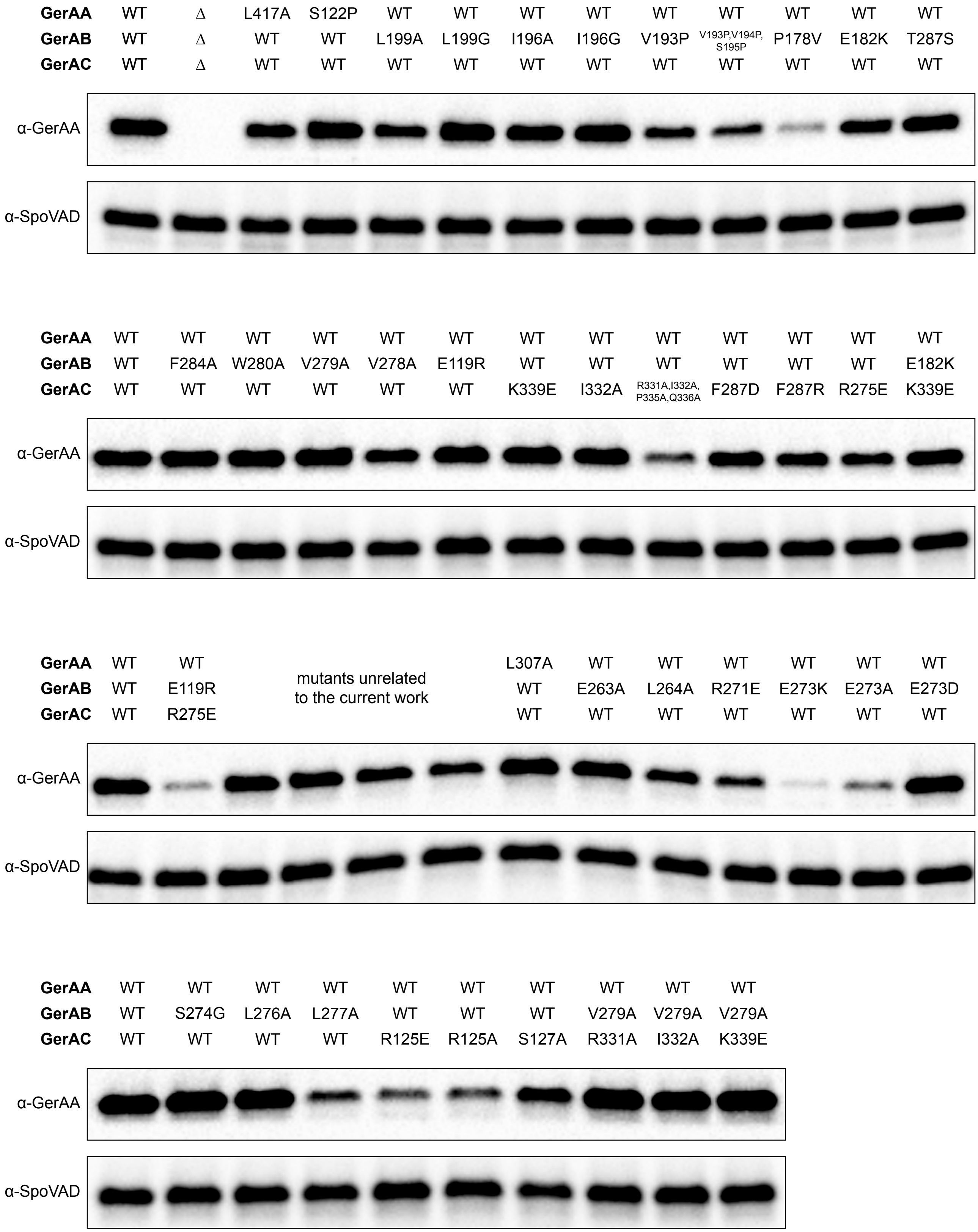


**Fig. S6.** Immunoblots assessing GerAA levels in the spores from **Fig. S5**. SpoVAD, loading control. See **Supp. Note 1** for details of assay interpretation.

**A**

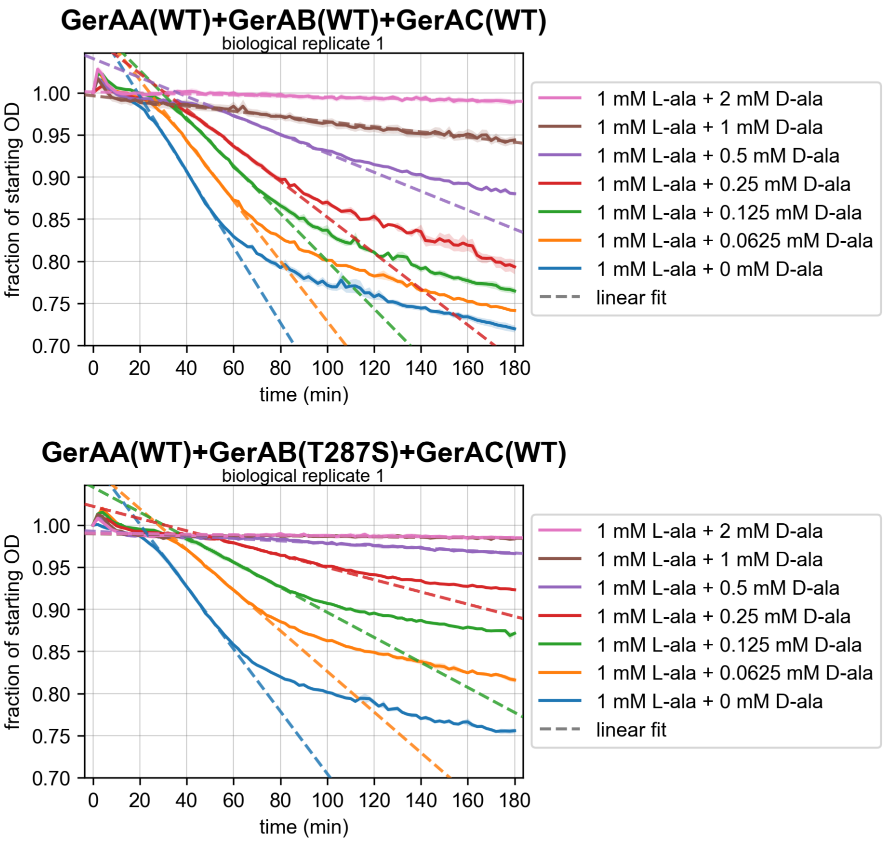


**B**

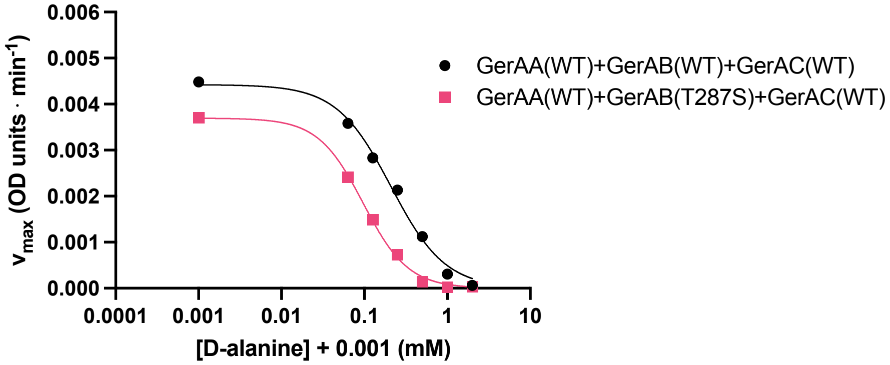


$$v_{max} = \frac{v_{max,0}}{1 + (\frac{[\text{D-ala}]}{IC_{50}})^h}$$

| | $v_{max,0}$ (OD units · min$^{-1}$) | $IC_{50}$ (mM) | h | $R^2$ |
|---|---|---|---|---|
| **biological replicate 1** | | | | |
| GerAA(WT)+GerAB(WT)+GerAC(WT) | 0.0044 (0.0040 to 0.0049) | 0.21 (0.16 to 0.28) | 1.3 (1.0 to 1.8) | 0.993 |
| GerAA(WT)+GerAB(T287S)+GerAC(WT) | 0.0037 (0.0035 to 0.0039) | 0.10 (0.09 to 0.11) | 1.6 (1.4 to 1.9) | 0.998 |
| **biological replicate 2** | | | | |
| GerAA(WT)+GerAB(WT)+GerAC(WT) | 0.0054 (0.0049 to 0.0059) | 0.18 (0.15 to 0.24) | 1.5 (1.2 to 2.0) | 0.994 |
| GerAA(WT)+GerAB(T287S)+GerAC(WT) | 0.0042 (0.0039 to 0.0044) | 0.09 (0.08 to 0.10) | 2.3 (1.8 to 2.8) | 0.998 |

**Fig. S7.** (**A**) Germination kinetics of spores with WT GerA and GerAA containing GerAB(T287S), constant [L-alanine] and titrated [D-alanine]. Shaded area, standard error of the mean for three technical replicates. For conditions whose final OD was >0.94, a line was fitted to the full time course. Linear fits for all other conditions are described in **Supp. Note 1**. (**B**) The $v_{max}$ from each of the fitted models in (A) (sign flipped so that a more positive number indicates a faster rate of germination) was plotted against [D-alanine]. Results of fitting the depicted model to these data are displayed in the table. 95% confidence interval is in parentheses.

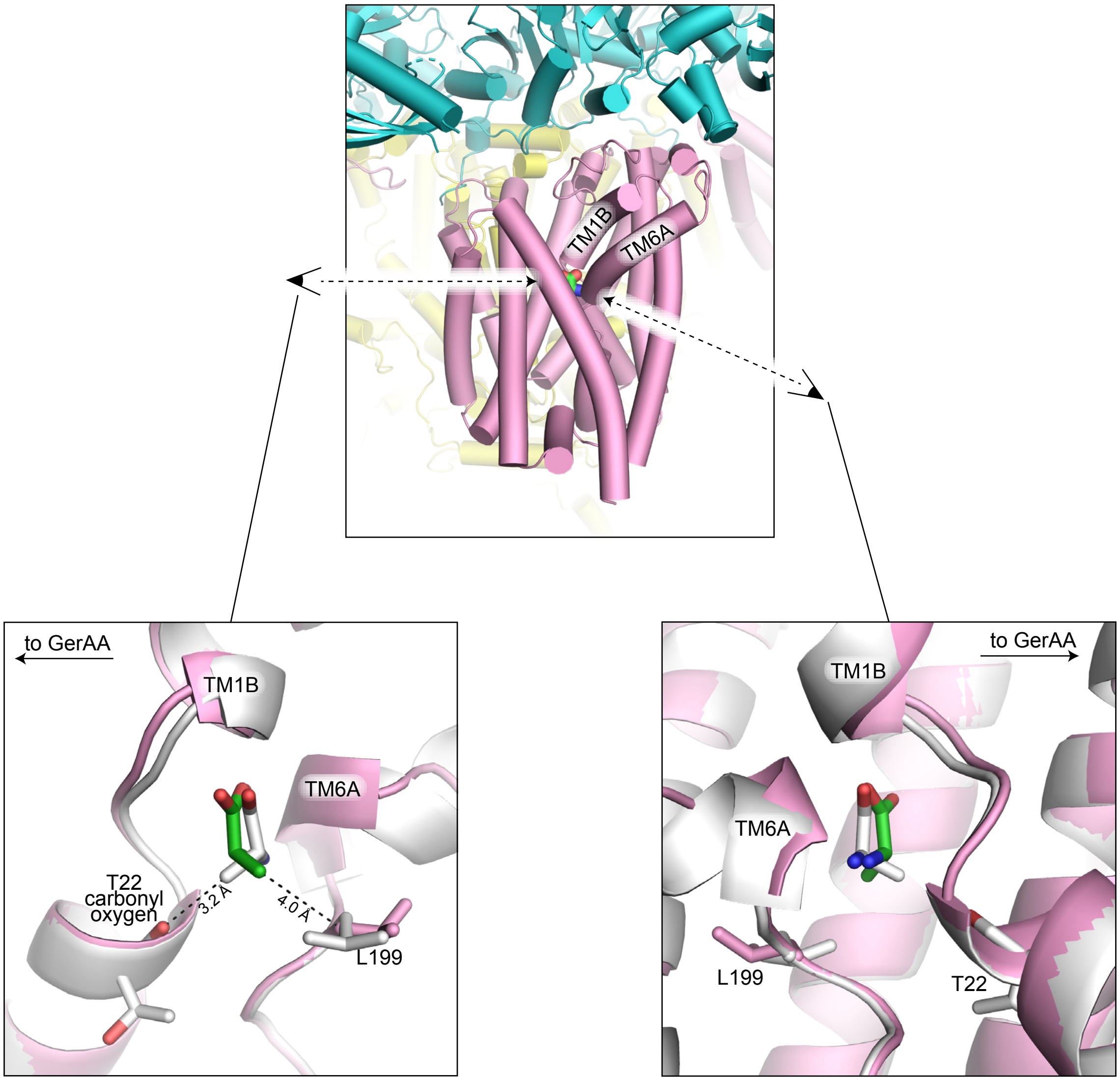


**Fig. S8.** Comparing the position of D-alanine (white, associated protein is white) and L-alanine (green, associated protein is pink) within the GerAB ligand-binding site of GerAA(A318C):GerAB:GerAC(S56C). See **Supp. Note 2**.

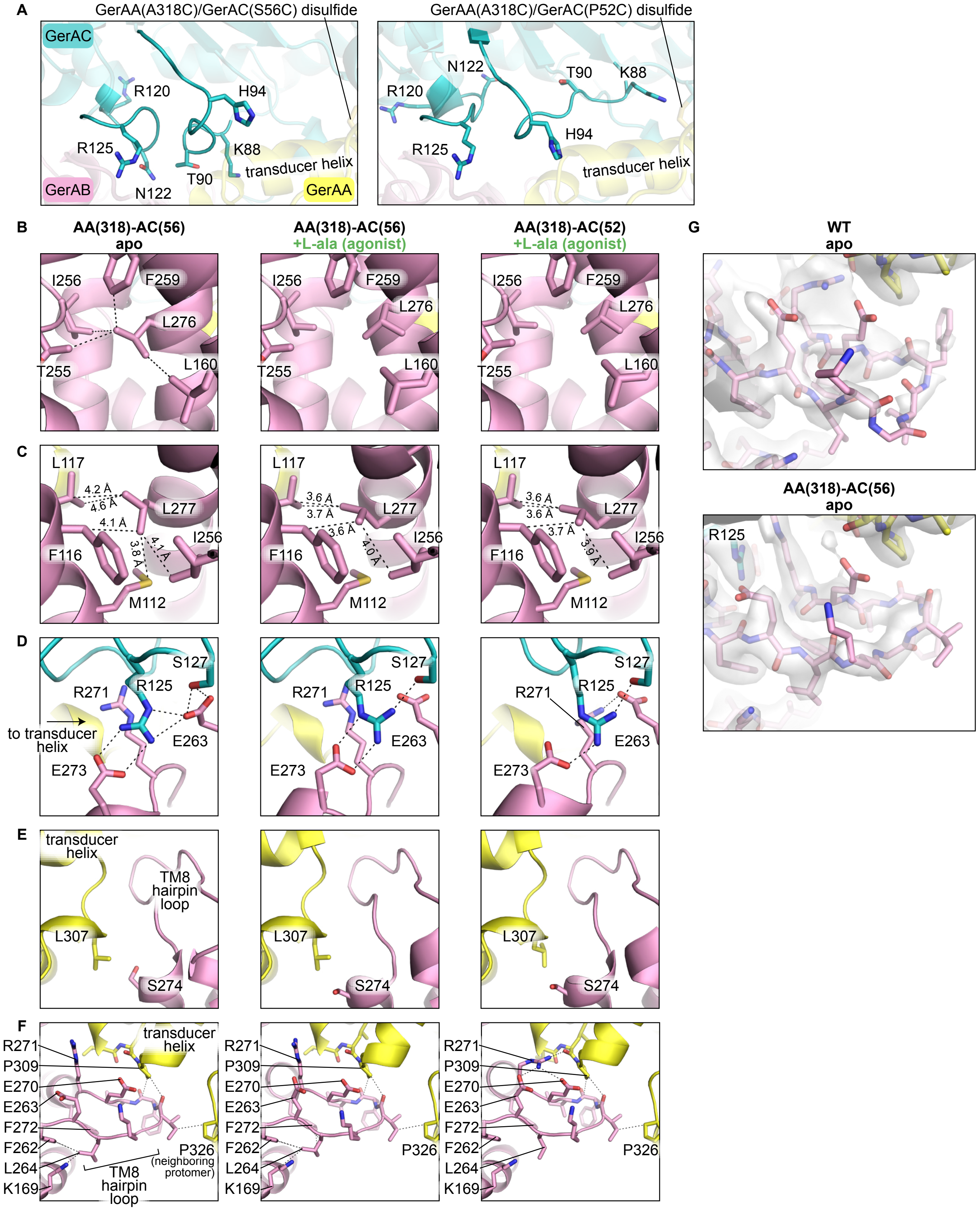


**Fig. S9.** See **Supp. Note 3** for discussion. (**A**) Loops in GerAC that pack differently in GerAA(A318C):GerAB:GerAC(S56C) vs. GerAA(A318C):GerAB:GerAC(P62C). Both depicted structures are L-alanine-bound. (**B**-**F**) Details of structural features that control Switch 2 propagation. GerAB-I256 is modeled with two alternative rotamers. (**G**) Cryo-EM maps (composite, unsharpened) and models demonstrating GerAC-induced ordering of the GerAB-TM8 hairpin loop. Compare to (F) for residue identification.

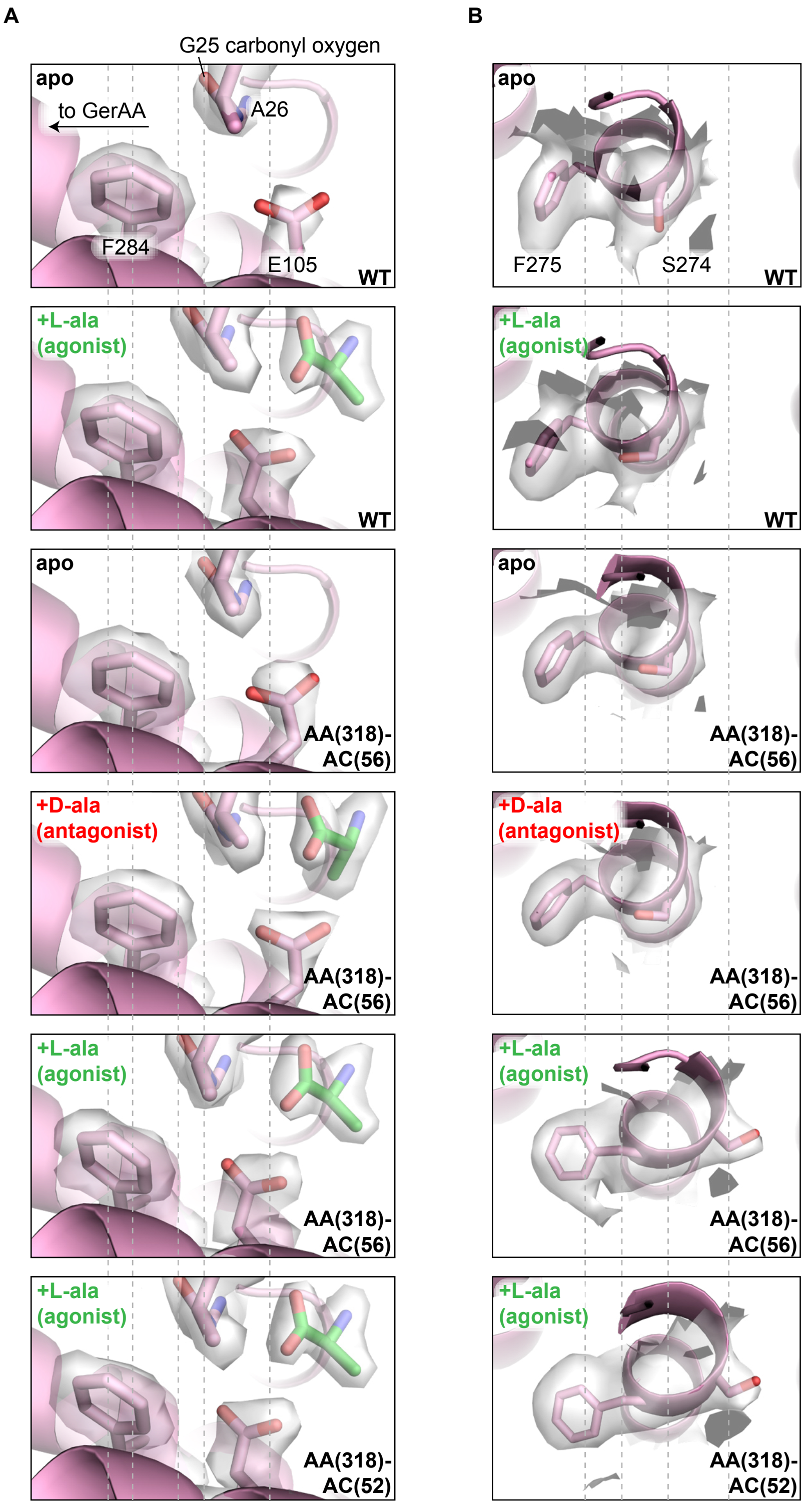


**Fig. S10.** (**A**) Cryo-EM maps (GerAB-focused, sharpened) and models demonstrating ligand- and GerAC-dependence of full Switch 2 activation near the ligand-binding site. (**B**) Cryo-EM maps (consensus, unsharpened) and models demonstrating ligand- and GerAC-dependence of full Switch 2 activation at the top of TM8. Vertical dashed lines are to guide visual comparison of key structural features.

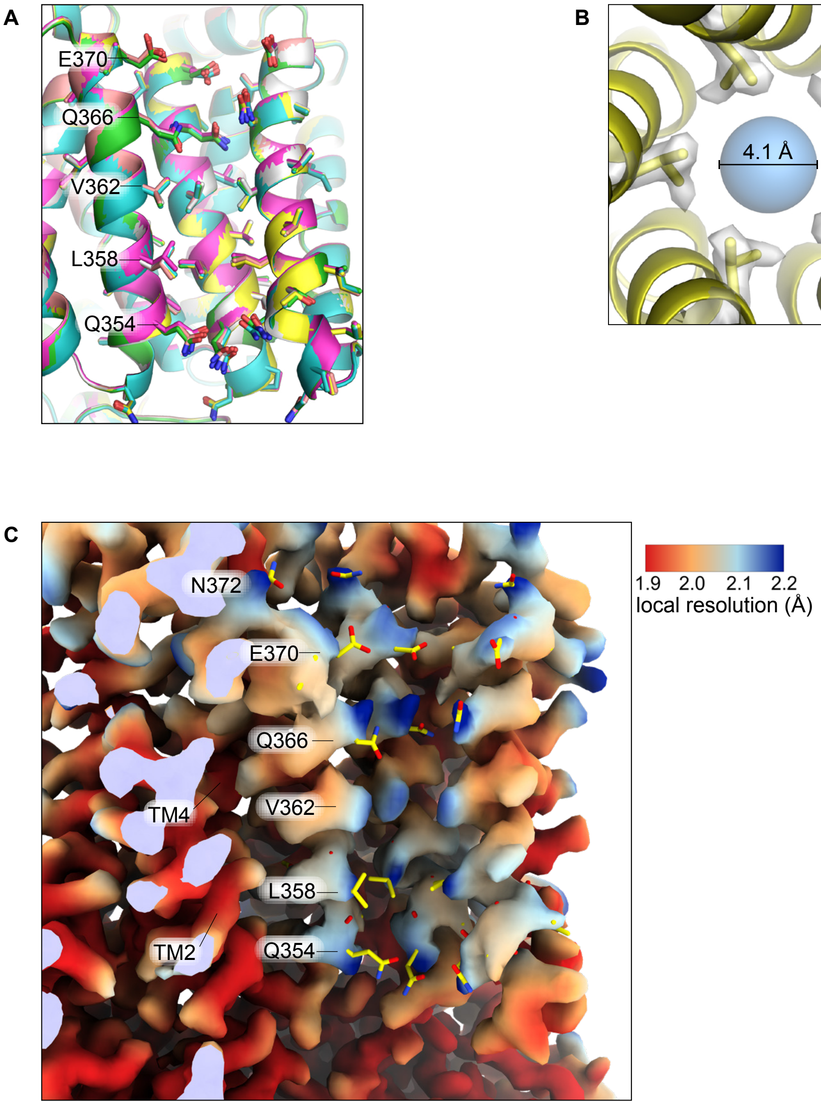


**Fig. S11.** (**A**) GerAA channel pore, equivalent view to **Fig. 4D**. Green, apo WT. Cyan, L-alanine-bound WT. Magenta, L-alanine-bound GerAA(A318C):GerAB:GerAC(P52C). White, apo GerAA(A318C):GerAB:GerAC(S56C). Salmon, D-alanine-bound GerAA(A318C):GerAB:GerAC(S56C). Yellow, L-alanine-bound GerAA(A318C):GerAB:GerAC(S56C). (**B**) Cryo-EM map (consensus, sharpened) and model at pore-facing residue V362 of L-alanine-bound GerAA(A318C):GerAB:GerAC(S56C). The blue sphere represents the MOLE-calculated (*66*) pore diameter at V362, based on the van der Waals surface of the carbon atoms (see Methods). This is an upper limit on the true pore diameter; modeling of hydrogen atoms would constrict the pore further. (**C**) Cryo-EM map (consensus, unsharpened, colored by local resolution) and model (yellow) depicting the same three GerAA protomers of L-alanine-bound GerAA(A318C):GerAB:GerAC(S56C) depicted in (A), along with surrounding helices. Note that because V362 is well-ordered, its model atoms are contained within the volume surface and therefore invisible. In contrast, the model atoms of other pore-facing sidechains exit the volume surface, reflecting their exploration of several different rotamers and positions.

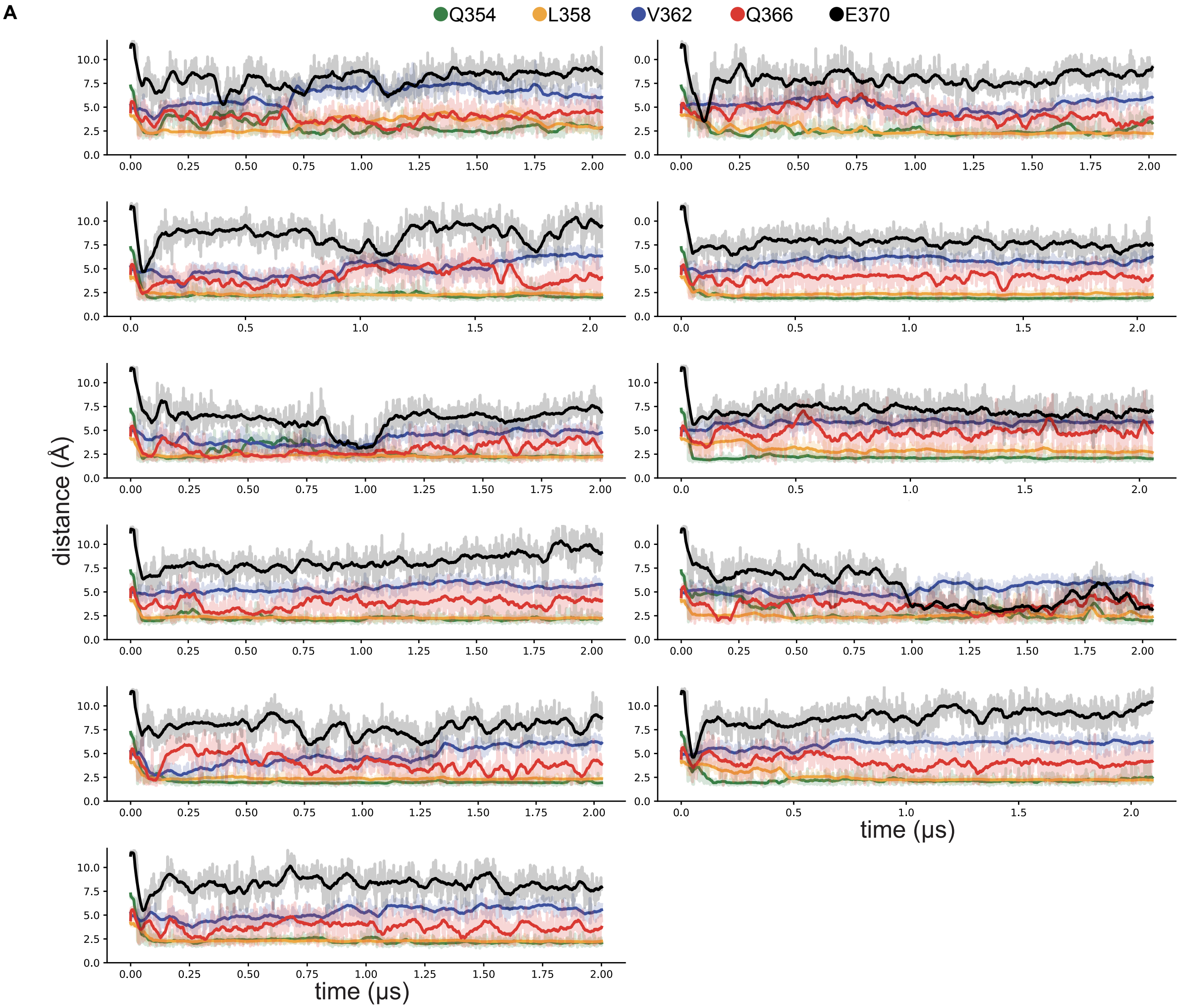


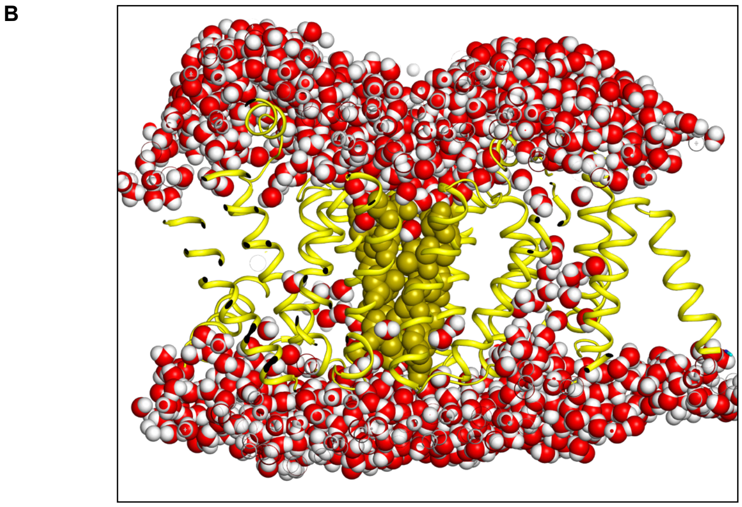


**Fig. S12.** (**A**) Additional independent MD simulations. See **Fig. 4A**. (**B**) Frame from equilibrated MD simulation. Yellow ribbon, GerAA transmembrane domain. Yellow spheres, pore-facing residues. Red/white spheres, water molecules. The inaccessibility of the pore to water molecules supports the assignment of this channel conformation as closed (*30*).

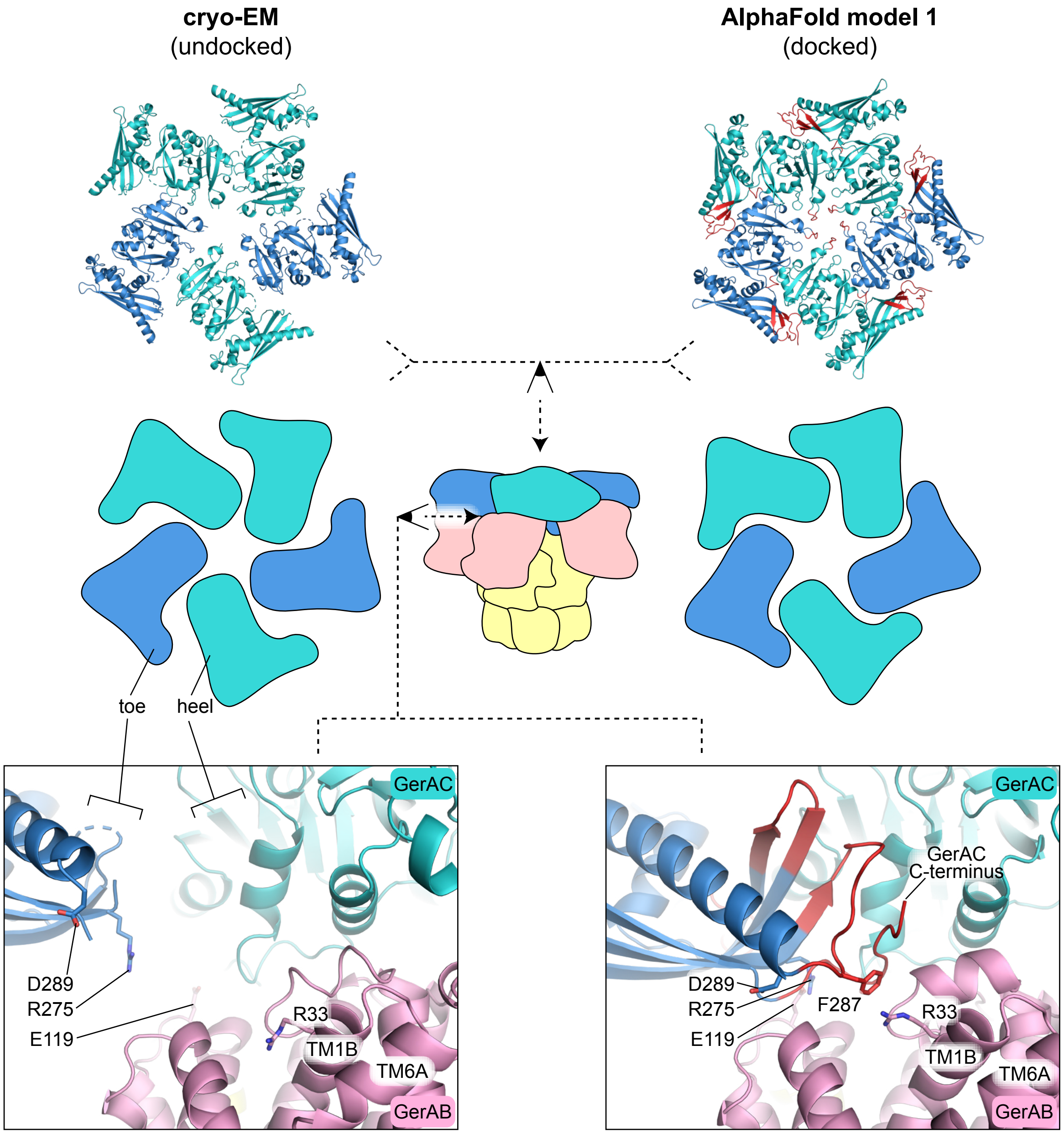


**Fig. S13.** GerAC toe-heel docking. See **Supp. Note 4**. The depicted cryo-EM model (L-alanine-bound GerAA(A318C):GerAB:GerAC(S56C)) is a representative of all experimental structures, which all exhibited a similar undocked conformation. Two GerAC protomers are colored a darker shade of blue to help visually distinguish which pieces of the model belong to which protomer. Amino acid residues that are present in the AlphaFold model but not the cryo-EM model (due to disorder) are colored red. Residues shown in stick representation are discussed in **Supp. Note 4** except for D289, which is displayed to help visualize how far the ordered part of GerAC travels between the undocked and docked states.

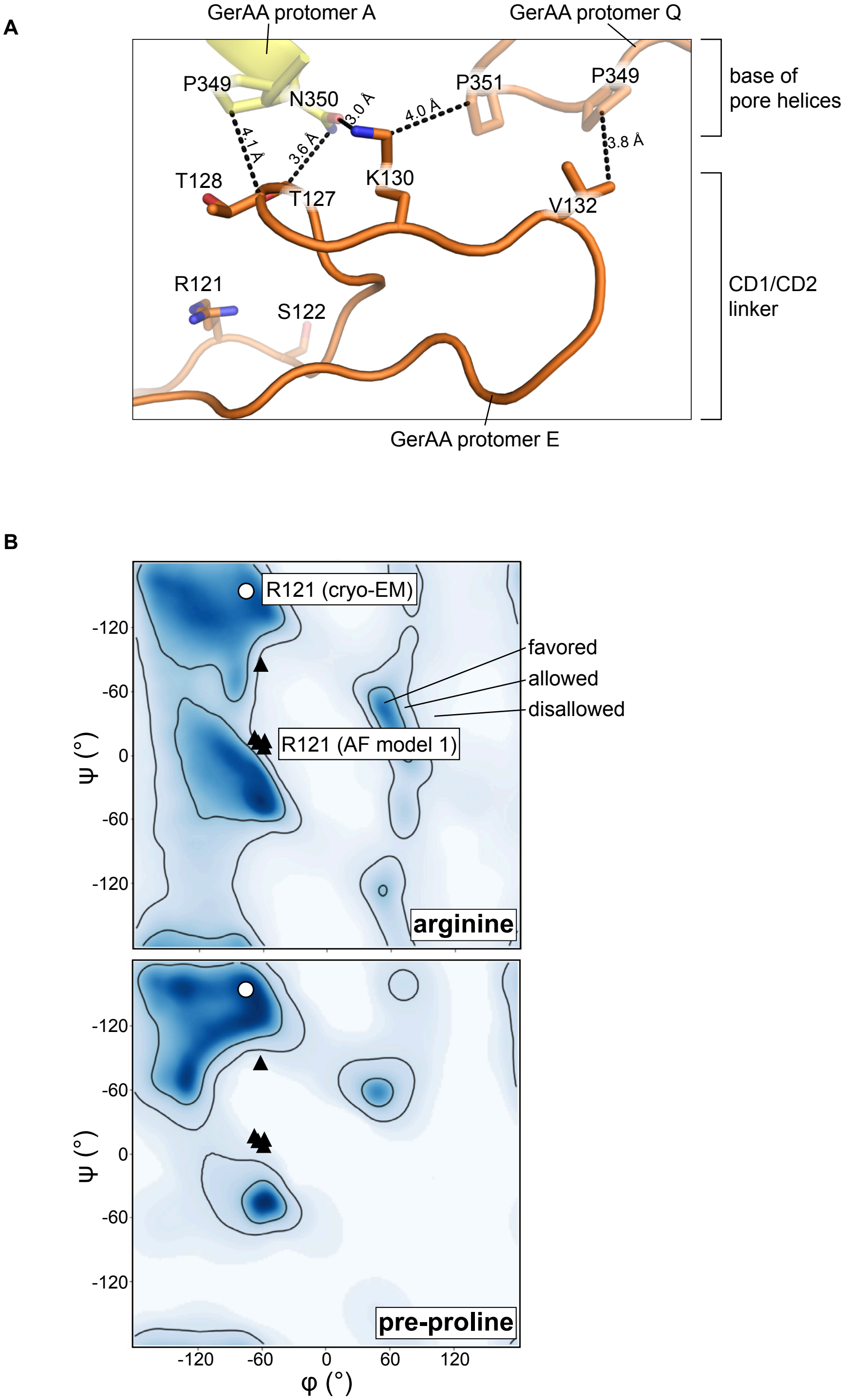


**Fig. S14.** (**A**) Detail of the central panel in **Fig. 4B**, contact between the CD1/CD2 linker and the base of the pore helices in GerAA. GerAA protomer IDs (A, Q, E) correspond to the chain IDs in the PDB file. (**B**) Ramachandran plots (from Phenix) for an arginine residue in non-pre-proline (top) vs. pre-proline (bottom) positions. Backbone dihedral angles of GerAA-R121 in the cryo-EM (white circle) and AlphaFold (black triangle) models are superimposed. Only one white circle is visible because the cryo-EM model has perfect $C_5$ symmetry at this position (see Methods). Note that a proline at position 122 would place the AlphaFold model (coaxial) but not the cryo-EM model (staggered) in the disallowed region.

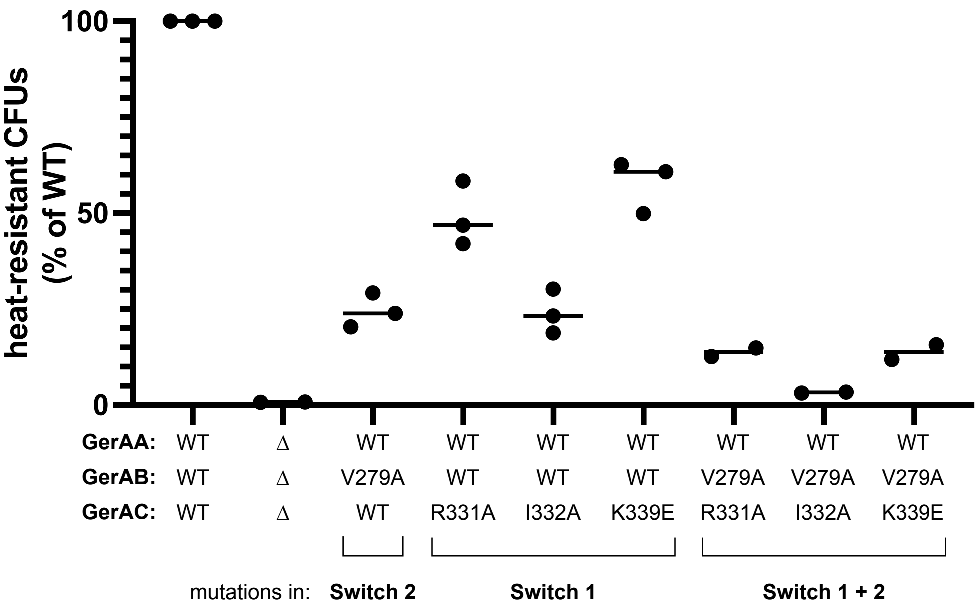


**Fig. S15.** Quantification of heat-resistant spore formation by *B. subtilis* containing mutations in either Switch 1 alone, Switch 2 alone, or both Switch 1 and Switch 2.

**Table S1. (separate file)**
Cryo-EM data collection, data processing, and model refinement.

**Table S2. (separate file)**
Local conformational signatures of AlphaFold predictions.

**Table S3. (separate file)**
*Bacillus subtilis* strains.

**Table S4. (separate file)**
Plasmids.

**Data S1. (separate file)**
Plasmid sequences.

**Data S2. (separate file)**
AMBER-relaxed AlphaFold predictions.
